# Deficiency of *ZC3HC1* modulates vascular smooth muscle cell phenotype and increases neointima formation

**DOI:** 10.1101/2021.09.29.462212

**Authors:** Redouane Aherrahrou, Tobias Reinberger, Julia Werner, Miriam Otto, Jaafar Al-Hasani, Maria Loreto Munoz-Venegas, Xiaoke Yin, Manuel Mayr, Mete Civelek, Heribert Schunkert, Thorsten Kessler, Jeanette Erdmann, Zouhair Aherrahrou

## Abstract

**Rationale:** The *ZC3HC1* gene has been linked to various cardiovascular traits. One variant, rs11556924-T, has been found to lower the risk of coronary artery disease (CAD) and blood pressure but increases carotid intima-media thickness (IMT).

**Objective:** This study aimed to determine how ZC3HC1 affects IMT using *in vitro* and *in vivo* models.

We analyzed the effect of the rs11556924-T allele on *ZC3HC1* expression in vascular smooth muscle cells (SMCs) from 151 multi-ethnic heart transplant donors. The results showed that rs11556924-T was associated with lower *ZC3HC1* expression and faster SMC migration. *ZC3HC1* knockdown (KD) experiments supported these findings, showing increased migration and proliferation. Mechanistically *ZC3HC1* KD led to decreased expression of contractile marker genes and the accumulation of cyclin B1, a key cell cycle protein. Pathway analysis of differentially expressed genes between *ZC3HC1* KD and controls SMCs showed decreased expression of genes in the cell division and cytoskeleton organization pathways, as well as higher expression of genes involved in extracellular matrix organization and cytokine-mediated signaling.

To validate these findings *in vivo*, we generated and characterized knockout (*Zc3hc1^-/-^*) mice. These mice had enhanced neointima formation in response to arterial injury and faster SMCs migration ability. However, complete loss of *Zc3hc1* led to a significant reduction in SMC proliferation and lower cyclin B1 protein level. In addition, immunostaining and confocal microscopy demonstrated, for the first time, that ZC3HC1 and Cyclin B1 were located at the cleavage furrow during mitotic progression of SMCs.

**Conclusions:** Collectively, our study suggests that lower ZC3HC1/NIPA level leads to increased SMC migration and neointima formation. Moreover, we proposed a biphasic role of NIPA in proliferation. Lower levels of NIPA promote SMC proliferation, while complete loss of NIPA hampers cell division and abrogates proliferation.

## Introduction

Coronary artery disease (CAD) caused by atherosclerosis is the leading cause of death in the developed world ^1^. Due to atherosclerotic deposits, the lumen of the vessels becomes narrow, limiting the supply of oxygen-rich blood to the affected tissue. This reduction in blood flow may result clinically in the development of chest pain and, in the event of atherothrombosis, in myocardial infarction (MI). Implantation of coronary stents is the primary therapeutic option to reopen an occluded coronary artery in MI patients. Despite the development of drug-eluting stents and balloons, restenosis due to neointima formation remains a major limitation after coronary intervention.

Genome-wide association studies (GWAS) have identified the single nucleotide polymorphism (SNP) rs11556924 at chromosome 7q32.2 CAD susceptibility locus, which harbors the *ZC3HC1* gene (Zinc Finger C3HC-type Containing 1; HGNC:29913) ^2–8^. The CAD-associated variant rs11556924-T is a non-synonymous variant, leading to an amino acid substitution p.Arg363His in the canonical transcript of *ZC3HC1* and a ∼4.9% to 7.0% reduction in CAD risk per allele ^4,6,9,10^. However, the same genetic variant increases the risk of carotid intima-media thickness (IMT) in patients with rheumatoid arthritis ^11^.

*ZC3HC1* encodes the nuclear-derived protein, nuclear interaction partner of anaplastic lymphoma kinase (NIPA)^12^, an essential component of the SCF-type E3 ubiquitin ligase complex that initiates the degradation of cyclin B1 ^7,13^. Because cyclin B1 and cyclin-dependent kinase (Cdk) form the M phase promoting factor, NIPA is involved in regulating the cell cycle at the G2 to M phase transition ^13^. In addition, the zinc finger protein ZC3HC1/NIPA has been recently referred to as a genuine nuclear basket protein of 53–55 kDa in vertebrates (∼84% similarity to the murine ortholog^14^) residing in the nuclear envelope and interacting with TRP (Translocated Promoter Region, Nuclear Basket Protein) encoding coiled-coil proteins ^15^ which are involved in mRNA export, chromatin compartmentalization in the interphase, and mitotic spindle checkpoint signaling during mitosis ^16^.

To understand the link between NIPA modulation and vascular remodeling, we assessed the role of *ZC3HC1* in human and murine vascular smooth muscle cells (SMCs), a key cell type involved in neointima formation following injury ^17^. Specifically, we analyzed the effects of *ZC3HC1* on migration and proliferation of primary human and mouse aortic SMCs *in vitro*. Subsequently, we used wire injury to induce neointimal hyperplasia ^18^ *in vivo* in *Zc3hc1*-deficient mice. Our findings showed that *ZC3HC1* deficiency was accompanied by cyclin B1 accumulation, increased migration, modified proliferation rates of vascular SMCs and enhanced neointima formation in mice.

## Material and Methods

### Expression quantitative trait locus analysis of ZC3HC1 in human SMCs

Primary human aortic SMCs isolated from the ascending aortas of 151 heart transplant donors at UCLA or obtained from commercial suppliers (Lonza and PromoCell) as this has been previously described ^19^. The University of Virginia IRB (Institutional Review Board) has ruled that specimens used for cell collection do not fall under the purview of current regulations governing the participation of human subjects in research since these specimens have no identifying information. All SMCs were maintained in Smooth muscle cell Basal Medium (SmBM, CC-3181, Lonza) supplemented with Smooth muscle Medium-2 (SmGM-2, CC-4149, Lonza). Briefly, the stranded libraries of high-quality ribosomal RNA-depleted total RNA were sequenced to ∼100 million read depth with 150 bp paired-end reads at the Psomogen sequencing facility. The reads with average Phred scores <20 were trimmed using Trim Galore, followed by mapping the reads to the hg38 version of the human reference genome using the STAR Aligner^20^ in two-pass mode. Gene expression of *ZC3HC1* was quantified by calculating the number of transcripts per million (TPM) using RNA-SeQC^21^. Expression quantitative traits locus (eQTL) analysis was performed using tensorQTL^22^ after correcting for sex, genotype PCs, and hidden confounding variables.

### Quantification of migration in 151 primary SMCs

Cell migration assays were performed with the xCELLigence Biosensor System using specifically designed 16-well plates equipped with membranes having 8-μm pores (CIM-plate 16; Roche Diagnostics). Cells in serum-free medium were seeded in the upper chambers, and the chemoattractant PDGF-BB (100 ng/mL) added to the lower chambers, with serum-free medium being the negative control. Cell migration was monitored over 24 h. Data were analyzed using RTCA software version 1.2 (Acea Biosciences Inc., San Diego, CA) combined with R software. The migration rate is displayed as normalized Log10 of time needed to reach saturation in the xCELLigence Biosensor build-up curve. The association between the genotype of rs11556924 and migration was calculated using linear mixed model to account for multiethnic population composition as previously been described ^19^.

### Silencing of ZC3HC1 gene expression using siRNA

Dicer small interfering RNA (siRNA) targeting the *ZC3HC1* gene (*ZC3HC1* siRNA, IDT-ID: hs.Ri.ZC3HC1.13.2) and scramble control siRNA were purchased from Integrated DNA Technologies (IDT). Human aortic SMCs (Cell Applications, Inc., #354-05a, C/C genotype for rs11556924) in M231 cell culture medium (Gibco) with Smooth Muscle Growth Supplement (SMGS) (Gibco) were cultured in 48-well cell culture plates (Greiner bio-one) at a density of 0.4 × 10^5^ cells per well for 24 h. The cells were transfected with 5 to 30 nM *ZC3HC1* siRNA or 5 to 30 nM control siRNA overnight using GenMute™ SMC siRNA Transfection Reagent (SignaGen Laboratories) according to the manufacturer’s instructions. The transfected cells were subsequently incubated in M231 SMGS culture medium for 48 h, after which the samples were harvested and stored at −80°C until used for qPCR and Western blot analyses.

### Cell migration and proliferation of ZC3HC1-knockdown SMCs

To determine the role of *ZC3HC1* gene expression knockdown (KD) in SMC migration, wound-healing assays with ibidi 4-well culture inserts in a 12-well plate format were performed as described ^23^. Briefly, approximately 2.2 × 10^4^ SMCs (Cell Applications, Inc., #354-05a, C/C genotype for rs11556924) per well were seeded into insert wells and incubated at 37°C and 5% CO_2_ for 24 h. Following siRNA transfection overnight (see above), the cells were incubated in M231 culture medium supplemented with Smooth Muscle Differentiation Supplement (SMDS) (Gibco), which contains only 1% (v/v) fetal bovine serum (FBS) and 30 µg/mL heparin. After 48 h, the inserts were removed, and the cells were washed with phosphate buffered saline (PBS) and cultured in M231 SMDS medium supplemented with 5 ng/mL PDGF-BB (Peprotech) to provoke cell migration. Images taken at 0 and 12 h with an Olympus IX70 microscope were analyzed using an in-house Python script. In brief, images were converted into black-and-white images and optimized using the methods *GaussianBlur* and *adaptiveThreshold* in the OpenCV package *cv2*. The mean pixel distances were calculated relative to time 0. All experiments were performed in triplicate. To assess the proliferation of *ZC3HC1* KD SMCs, the cells were plated into 96-well plates (0.6 × 10^4^ cells/well) and transfected as described above. The next day, the cell culture medium was replaced by M231 medium supplemented with 1% FBS to starve the cells. These human SMCs were subsequently treated with 100 ng/mL PDGF-BB (Peprotech) to induce proliferation. To quantify the proliferation rate, the cell nuclei were stained with Hoechst 33342 dye (ThermoFisher Scientific) at several time points and the number counted using an in-house Python script *CellCounter.py*. Six wells were analyzed per condition, with all experiments performed in triplicate. The values were normalized to time point zero to account for small differences in initial cell numbers after siRNA transfection. To confirm the proliferation results, a Bromodeoxyuridine (BrdU) based proliferation assay was conducted according to the manufacturer’s introductions (Roche, # 11647229001). The incorporation of BrdU was detected calorimetrically after six to eight hours of incubation time at a wavelength of 370 nm and normalized to cells devoid of BrdU. In addition, cell nuclei were stained with Hoechst 33342 dye. Proliferation assays were performed in three and five replicates, respectively.

### qPCR analysis

These analyses were performed as described ^21^ ^22^. Briefly, total RNA was isolated from cultured cells using RNeasy plus kits (Qiagen, Valencia, CA, USA) and reverse transcribed into cDNA. mRNA levels were determined by relative quantitative RT-PCR and analyzed using the ΔΔCT method relative to the internal standard, GAPDH/Gapdh ^24^. The primers (Eurofins Genomics) used in this study are shown in **Table 1**.

**Table 1:**
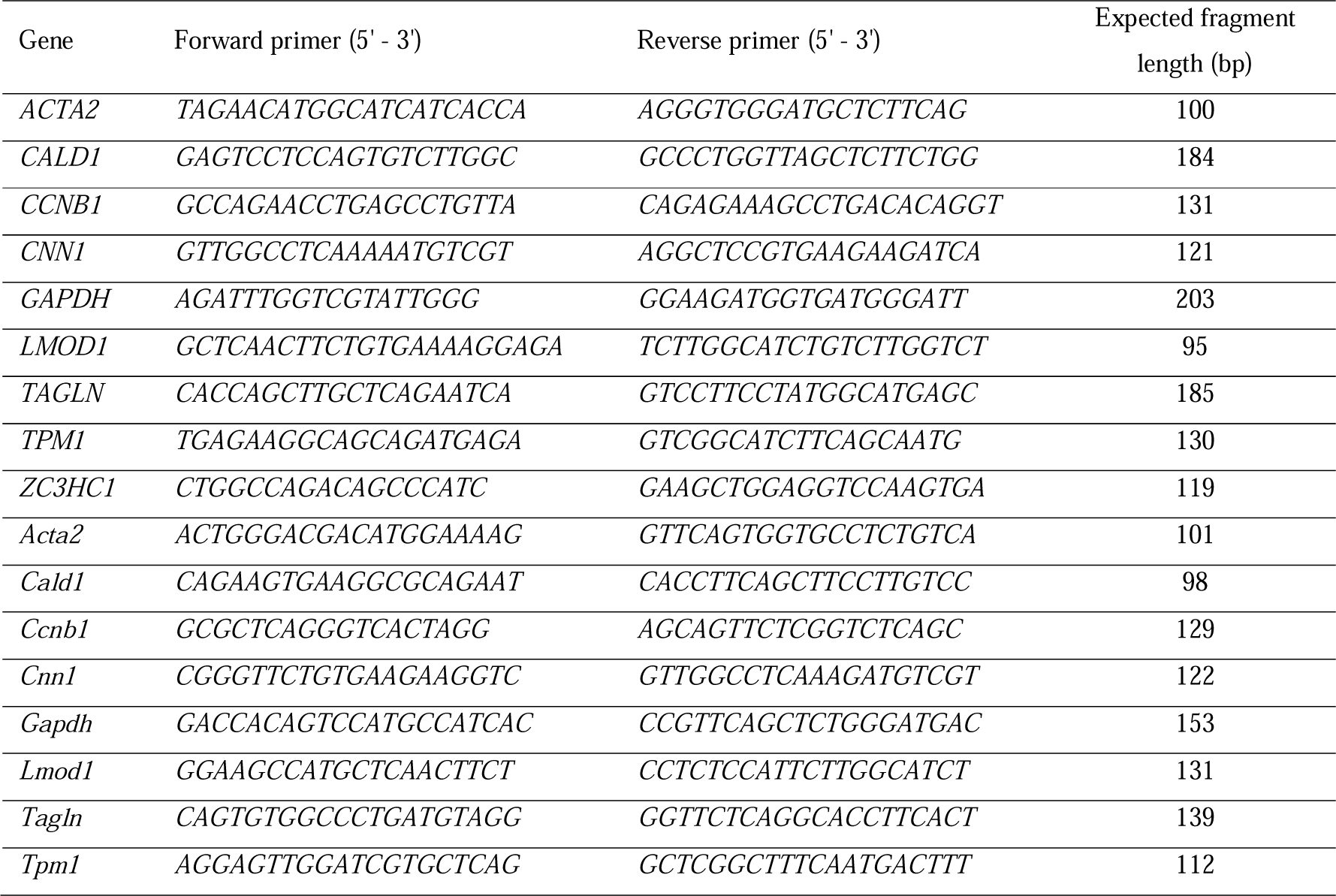
Primer sequences used in qPCR reactions. Human genes are in upper case, while the mouse genes are in lower case.

### Western blot analysis

Western blot analyses were performed as previously described ^25^. Briefly, 15 to 75 µg samples of protein isolated from cultured cells were electrophoresed on SDS-PAGE gels and transblotted onto nitrocellulose membranes. The blots were treated with 5% skim milk and incubated with primary antibodies, including anti-NIPA (phospho S354) antibody (abcam ab63557), anti-cyclin B1 antibody (abcam ab181593), anti-SRF (Cell Signaling Technology, SRF (D71A9) XP), anti-CNN1 (abcam ab46794), anti-LMOD1 (Proteintech 15117-1-AP), and anti-GAPDH antibody (loading control; abcam ab181602). Amido black dye (Sigma-Aldrich A8181-1EA) served as internal loading control. The blots were subsequently incubated with the appropriate secondary antibodies. Protein bands were detected using the ECL Prime Western Blotting Detection Reagent (GE Health Care) and quantified using ImageJ software.

### RNA sequencing

Human aortic SMCs (Cell Applications, Inc., #354-05a) were seeded into 6-well plates at a density of 3.1 × 10^4^ cells/cm^2^ and transfected with siRNA as described above. After 2 days in M231 SMDS medium, 5 ng/mL PDGF-BB (Peprotech) was added. Cells devoid of PDGF-BB served as an internal control. After incubation for 24 h, the RNA was extracted using innuPREP RNA Mini Kits 2.0 (Analytik Jena), yielding ∼5 µg total RNA (RIN>7) per sample. RNA sequencing (RNAseq), including ribosomal RNA-depletion and quality control, was performed at the Novogene sequencing facility (NovaSeq 6000 PE150, 150 bp paired-end reads). After trimming reads with low average Phred scores (<20) using Trim Galore, all included samples passed the quality control analysis. Reads were mapped to the hg38 version of the human reference genome using the STAR Aligner^20^ in two-pass mode, with gene expression quantified by calculating TPM using RNA-SeQC^21^.

### Differential gene expression and functional enrichment analysis

A total of 17,242 genes with >6 reads were included in at least 20% of the samples in at least one of the two conditions (ZC3CH1 siRNA and control siRNA) for differential expression analysis using DESeq2 controlling for batch effect ^26^. Genes were considered to be differentially expressed under treated and untreated conditions when p_adj_ was <0.05 and log_2_(fold-change) was >0.4. Principal component analysis (PCA) was performed using R. PDGF-BB treatment was defined as a covariate. Network analysis and Gene Ontology (GO) enrichment after clustering were performed to characterize the functional consequences of differences in gene expression associated with downregulation and normal expression of *ZC3HC1* ^27^. In brief, a functional gene network containing differentially expressed genes (p_adj_<0.05) was constructed with the help of the STRING database repository (https://string-db.org/) using the REST API implemented in Python. The following parameters were used: species: 9606, network_type: functional, network_flavor: confidence, score>0.4, add_white_nodes: 30 for the complete gene network (**Supplementary Figure S2**) and 8 for the CCNB1 subnetwork (**Figure 3**). Agglomerative ward clustering of the gene network was performed using the *scikit-learn* (version 0.24.2) clustering package. Gene set enrichment of each cluster was performed using the REST API of STRING^27^. Gene networks were visualized by the Python packages *networkx* (V4.4.0) and *matplotlib* (V40.8.0).

### Proteomics

The proteomic analysis was conducted as described^28^. In brief, SMCs were transfected (30 nM siRNA) as described above. After incubation for three days in basic SMC medium, the cells were stimulated with 5 ng PDGF-BB for 24 h in M231 medium supplemented with 1% FBS. Primary SMCs from heart transplant donors carrying the T allele (n=7) or C allele (n=6) were cultured as described above. For both SMC types (knockdown/control and heart transplants), cells were washed three times with cold phosphate-buffered saline (PBS) thoroughly, detached, and collected using centrifugation. After the addition of cell lysis buffer (#9803, Cell signaling), including protease inhibitors/phosphatase inhibitors (#04906837001, ROCHE diagnostics/ #11697498001, ROCHE diagnostics), cells were sonicated on ice two times for 15 seconds and left on ice for 20 min. The supernatant was subjected to mass spectrometry (MS)^28^. Normalized abundance values were kept if at least two unique peptides per protein were detectable and more than 30% of samples exhibited MS signals for the corresponding protein. In a pairwise comparison, missing values were replaced with zeros if more than 90% of replicates of group A show signals and less than 10% of replicates of group B lacked MS signals. After log2 transformation, missing values were imputed using k-nearest neighbors imputation and differential protein levels were assessed with the aid of the limma R package^29^, adjusting for multiple testing using the Benjamini-Hochberg method.

### Generation and housing of animals

Heterozygous *Zc3hc1* (^+^/^-^) mice (Zc3hc1tm1a(KOMP)Wtsi, background: C57BL/6N) were obtained from KOMP at UC Davis, California. They were created using a targeting construct of the mouse *Zc3hc1* gene, designed by introducing the EN2 splicing acceptor (EN2 SA) followed by the beta-galactosidase sequence and a neomycin selection cassette 3’ at exon 4. Heterozygous mice were initially backcrossed to a C57BL/6J genetic background for at least six generations and then used in Het × Het mating to generate sufficient numbers of *Zc3hc1* knockout (KO or ^-/-^) and wild-type (WT or ^+/+^) littermates for the experiments. Homozygous *Zc3hc1* (^-/-^) were found to be infertile. Mice were genotyped using PCR screenings of DNA samples isolated from ear biopsies. Mice were genotyped using the primers 5’-TTGACTGACAGAGGATGAGAGC-3’ (forward) and 5’-GGGCCTTTAATCCCAACACT-3’ (reverse), targeting the second *LoxP* site located between exons five and six in the *Zc3hc1* gene. The expected lengths of the DNA fragments were 298 bp for the knockout and 260 bp for the WT mice (**Figure 4A, right**). In addition, *β*-gal-activity was measured in heart cryosections from one mouse per genotype (*Zc3hc1* ^+^/^+^, or ^−/−^) (**Supplementary Figure S6C**), as described ^30^.

### Neointima formation mouse model

All animal experiments were locally approved by the German animal studies committee of Upper Bavaria and conformed to the guidelines from Directive 2010/63/EU of the European Parliament on the protection of animals used for scientific purposes. Wire injury was induced in 8- to 12-week-old female mice as described^31^ and established successfully in our group^32^. Briefly, all surgical procedures were performed under general anesthesia. Anesthesia was achieved by intraperitoneal (i.p.) injection of midazolam (5.0 mg/kg), medetomidine (0.5 mg/kg), and fentanyl (0.05 mg/kg) (MMF) in 300μl of 0.9% (w/v) sodium chloride. One-third of the dosage of MMF was added if the inter-toe reflex was re-established or the duration of anesthesia exceeded 60 minutes. After a skin incision, the left femoral artery was exposed by blunt dissection and an angioplasty guidewire (0.015 inches in diameter, No. C-SF-15-15, COOK, Bloomington, IN) was introduced into the arterial lumen and inserted towards the iliac artery via arteriotomy. The wire was left in place for 1 minute to denude and dilate the femoral artery. After that, the wire was removed and the blood flow of the femoral artery was restored. The arteriotomy site was ligated and the skin was closed. The antagonists atipamezole (2,5 mg/kg), flumazenil (0,5 mg/kg) and naloxone (1,2 mg/kg) were administered subcutaneously to terminate anesthesia. Mice were sacrificed 14 days later by i.p. injection of pentobarbital. To remove blood inside the femoral arteries, the mice were perfused via the left ventricle and over the descending aorta with PBS (pH= 7.4). Femoral arteries were harvested and fixed in 4% buffered paraformaldehyde overnight.

### Histology and morphology

Paraffin-embedded femoral arteries were sectioned at 2-µm intervals and stained with hematoxylin and eosin. For morphometric analysis, images were digitalized (Leica DFC450C camera) and serial sections in 50-µm intervals of each artery were blindly analyzed using ImageJ software (NIH Image Software). Media thickness (M) and the area of neointima formation (NI) were quantified and neointima-to-media ratios (NI/M) were calculated. Sections of paraffin-embedded femoral arteries were de-paraffined through a series of decreasing concentrations of alcohol (3x xylol, 2× 100% ethanol, 1× 96% ethanol, 1× 70% ethanol and deionized water). For staining, slides were stained using hematoxylin eosin (HE) solution (Carl Roth, #T865.1) or via. To fix the slides, they were treated with a series of increasing concentrations of alcohol (see above), followed by treatment with Cytoseal XYL (ThermoFisher Scientific #8312-4). For histochemistry staining, the slides were heated for 1.5 min in a microwave at 900 W and for 15 min at 90 W. For histochemistry, the slides were washed three times with PBS and unspecific binding was blocked by incubation with 5% (v/v) goat serum in PBS. Tissue slides were stained overnight at ambient temperature with anti-Ki-67 (1:100 in PBS, ThermoFisher Scientific, Clone:B56, #15898578), anti-ACTA2 (1:100 in PBS, Abcam ab5694), anti-LMOD1 (1:200 in PBS Proteintech 151117-1-AP), or anti-CD31 (1:50 in PBS, Abcam ab28364)antibody, and primary antibodies were visualized using a horseradish peroxidase system (Santa Cruz, SC2004) and DAB staining solution (DAKO, #3468). Cell nuclei were counterstained with hematoxylin. Slides were fixed aquatex (Merck #1085620050).

### Isolation and culture of murine SMCs

Mouse aortic SMCs were isolated from the thoracic aortas of WT and KO (*n*=8-10) as reported previously ^25^. Briefly, the thoracic aortas were collected, and fat and connective tissues were removed. The samples were pre-digested with collagenase II and the adventitia was removed mechanically. Adventitia-free aortas were cut into small pieces and further digested with collagenase II with shaking for at least 6 h, until complete tissue dissociation. The isolated cells were pelleted and resuspended in culture medium (DMEM plus 10% FBS and 1% penicillin/streptomycin, Gibco). Cells were expanded and grown on surfaces coated with 0.1% (w/v) gelatin. To quantify the purity of the populations, the isolated cells were characterized by flow cytometry using the SMC-specific FITC-labeled anti-*α*-SMA antibody (Sigma, F3777) and an isotype control (mouse IgG2a-FITC, Sigma F6522), yielding >95% purity (**Supplementary Figure S7**).

### Murine SMC migration and proliferation

The migration of murine aortic SMCs was assessed using the xCELLigence RTCA DP system as described above. Briefly, the electrode side of the membrane was coated with 0.1% gelatin. The bottom compartment of each well was filled with 165 μL of culture medium or serum-free medium as a control, and the membrane compartment was mounted. The upper wells were filled with 50 μL of serum-free medium and equilibrated for 1 h at 37 °C in an incubator. Cell concentrations were adjusted to 2.7 × 10^4^ cells/mL per well. Data were recorded for 24 h. To quantify the proliferation of murine aortic SMCs, cells were seeded in 12-well plates to yield a confluency of ∼30%. To make these results comparable to those of the proliferation assay for human SMCs transfected with siRNA, the cells were incubated for 1 day (with 100 ng/mL murine PDGF-BB or without), with the timepoint 0 h defined as 24 h after seeding. At each given timepoint, the cells were washed once with 1 mL PBS and fixed with 0.5 mL of 4% (v/v) paraformaldehyde (PFA) for 5 min. After removing the PFA, the cells were permeabilized with 0.5 mL Triton-X-100 for 10 min, washed with 1 mL PBS, and stained with 0.5 mL 4’,6-diamidino-2-phenylindole, dihydrochloride (DAPI) solution for 10 min in the dark. After acquiring images of 10 non-overlapping regions per well, both the cell number and average size of nuclei were determined using our in-house Python script *CellCounter.py*. The results were normalized to time point zero for each condition and replicate. The results were validated using the BrdU assay as described above. The incubation of BrdU was shortened to four hours. Proliferation assays were performed in five and four replicates, respectively.

### Immunofluorescence and confocal microscopy

SMCs were seeded on µ-slide chamber slides (ibidi) and cultured in SMC medium (see above) containing 100 ng/mL murine PDGF-BB until desired confluency. Both genotypes (Zc3hc1 KO vs. WT) were treated equally. After fixation with cold methanol-acetone solution, PDGF treated SMCs (see above) were permeabilized with 0.2% Triton-X (1% BSA in PBS) and unspecific sites were blocked with 3.5% BSA. SMCs were stained with anti-ZC3HC1 (Origene, AP20366PU-N, 1:100), anti-Tubulin (abcam, ab6160, 1:1000) or anti-cyclin B1 (Thermo Fisher Scientific, PA5-85113, 1:200), DAPI (4’,6-Diamidino-2-phenylindol), and Alexa Fluor 568/ Alexa Fluor 488. Images were acquired using a confocal microscope (Leica, TCS-SP5) and eight to ten z-stacks (∼1 µm distance) were combined for maximum intensity projections.

### Statistical Analysis

All data are presented as the mean ± standard deviation (SD), with two groups compared by unpaired Student’s t-tests or Mann–Whitney U tests (n<8 or not normally distributed data). All statistical analyses were performed using R or Python libraries, with p<0.05 considered statistically significant.

## Results

### Associations of the *ZC3HC1* variant rs11556924-T with cardiovascular disease (CVD)-related phenotypes

A hypothesis-free phenome scan was performed using the MR-Base PheWAS online tool^33^ and the GWAS Catalog^34^ to determine the associations of rs11556924 variants with CVD-related phenotypes. As expected, a genome-wide significant association was observed between this SNP and cardiovascular-related traits, including hypertension and CAD (**Figure 1A, Supplementary Table 1**), with the rs11556924-T allele associated with reduced risk of these diseases. A suggestive association (p=1.4 × 10^-5^) was also observed between this SNP and mean carotid IMT in the MRC-IEU consortium data (**Supplementary Table 1**).

**Figure 1.**
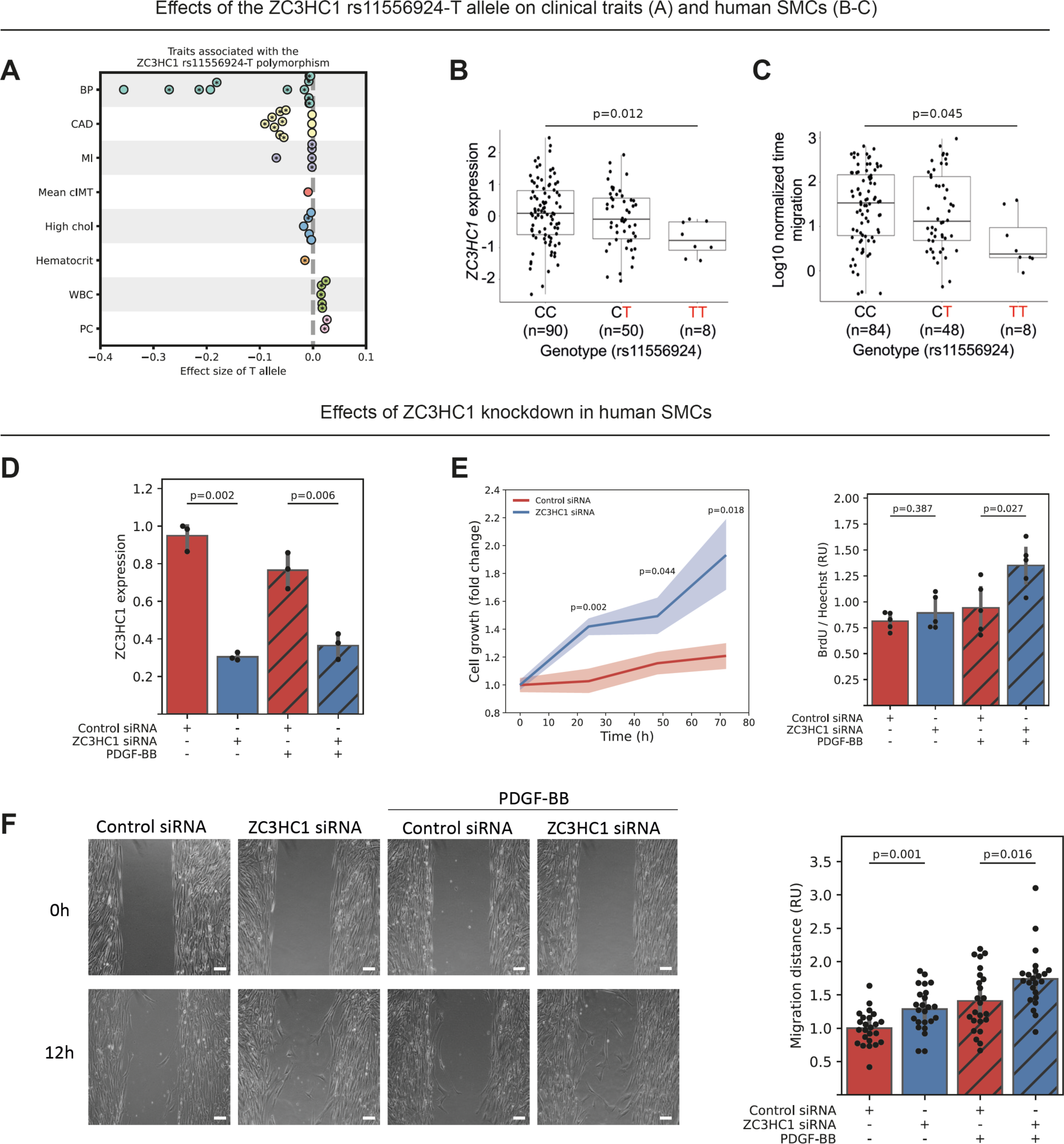
Association of rs11556924 with cardiovascular diseases predominantly driven by vascular SMC dysfunction, *ZC3HC1* expression and vascular SMC phenotype. **(A)** Predominant association of rs11556924 with cardiovascular disease (CVD) driven by vascular SMC dysfunction. The plot shows the beta values (effect size) of the rs11556924-T allele derived from several genome-wide association studies (GWAS) categorized according to eight CVD-related traits (BP, blood pressure; CAD, coronary artery disease; MI, myocardial infarction; cIMT, carotid intima-media thickness; chol, cholesterol; WBC, white blood cell count; PC, platelet count). Asterisks indicate genome-wide significance (p<5 × 10^-8^); data points without asterisks represent genome-wide associations with suggestive significance (p<5 × 10^-5^). Negative effect size indicates that the risk allele (T) of the single nucleotide polymorphism, rs11556924, is associated with a lower risk for several traits, including blood pressure or coronary artery disease. **(B)** The risk allele (T) of rs11556924 is associated with lower *ZC3HC1* expression (TPM, transcripts per million) and **C)** faster migration by vascular SMCs. SMCs transfected with siRNA against *ZC3HC1* showed significant **(D)** *ZC3HC1* downregulation, **(E)** higher PDGF-induced proliferation rate confirmed by counting nuclei stained with Hoechst 33342 dye (left, n=3, six images per replicate) as well as BrdU assay (right, n=5, mean of three to four wells per condition), and **(F)** increased migration in the presence or absence of PDGF-BB (n=3, eight images per replicate). Representative images of migratory SMCs transfected with *ZC3HC1* siRNA and control siRNA are shown after 12 h (scale bars, 100 µm). Values are shown as means. The translucent error bands represent one standard deviation.

### The rs11556924-T SNP is associated with lower *ZC3HC1* expression and higher migration in human aortic SMCs

To understand the role of the rs11556924 variant at the level of gene expression, an eQTL analysis was performed using aortic SMCs of 151 human donors. This assay showed that the T allele was associated (p=0.012) with lower *ZC3HC1* expression (**Figure 1B**). We observed 5% and 10% reduction of *ZC3HC1* expression in SMCs heterozygous and homozygous for the T risk-allele respectively, compared to the homozygous carriers of the C non-risk-allele. To our knowledge, there are no eQTLs in monocytes/macrophages^35^ or aortic endothelial cells^36^, suggesting that the regulatory impact of rs11556924 in the *ZC3HC1* locus is SMC specific. Therefore, we assessed the impact of this variant on SMC migration, a hallmark for vascular remodeling ^17^. Cell migration assays were performed using 100 ng/mL PDGF-BB as a chemoattractant in serum-free media, as described previously ^19^. SMCs carrying the rs11556924-T variant were found to migrate faster toward PDGF-BB than SMCs carrying the C allele (**Figure 1C**). The heterozygous and homozygous SMCs for the T risk allele migrate 10% and 50% faster respectively, compared to the homozygous carriers of the C non-risk-allele.

### Downregulation of *ZC3HC1* in human SMCs increases proliferation and migration

To further investigate how *ZC3HC1* modulation affects SMC proliferation and migration, cells were transfected with siRNA against *ZC3HC1* (*ZC3HC1* siRNA; knockdown efficiency was ∼70% at 12 h after transfection (**Figure 1D**)) or control siRNA. A lower level of *ZC3HC1* mRNA significantly enhanced the proliferation of SMCs (**Figure 1E**). Of note, no significant correlation was observed in SMCs derived from 151 human donors^19^ between the T allele and cell proliferation at 24 h. In agreement with the migratory response of the 151 SMC types, siRNA mediated knockdown of *ZC3HC1* promoted cell migration (**Figure 1F**). Moreover, adding PDGF-BB (mimicking vessel injury) was accompanied by lower expression of *ZC3HC1* and increased migration in control SMCs as expected (**Figure 1D-F**). An increase of migration rate was also observed in *ZC3HC1* KD compared to control SMCs in the presence of PDGF-BB.

### Transcriptional profiling of *ZC3HC1* downregulation in human SMCs

To determine the transcriptional profile of *ZC3HC1* downregulation, transcriptome analysis using RNAseq was performed in aortic SMCs in the presence or absence of 5 ng/mL PDGF-BB. After quality control and quantification, the number of expressed genes (defined as genes with more than six read counts in at least 20% of the samples) ranged from 16,880 under control conditions to 17,242 after treatment with PDGF-BB (**Supplementary Table 5**). PCA identified two distinct clusters of samples, corresponding to the cells cultured under the two conditions with endogenous *ZC3HC1* expression and *ZC3HC1* downregulation (**Supplementary Figure S1**). Further, 3,045 genes were differentially expressed (p_adj_<0.05). Of these, 284 genes showed a log_2_(fold-change [FC]) above 0.5 (median 0.68), and 179 genes were downregulated with log_2_(FC)<-0.5 (median −0.67). The latter also includes the canonical SMC markers *LMOD1*, *TPM1*, *CNN1*, *CALD1*, *ACTA2*, and *TAGLN* (**Figure 2A-B**). Downregulation of the expression of genes encoding SMC markers in response to *ZC3HC1* knockdown was confirmed by qPCR (**Figure 2C**).

**Figure 2.**
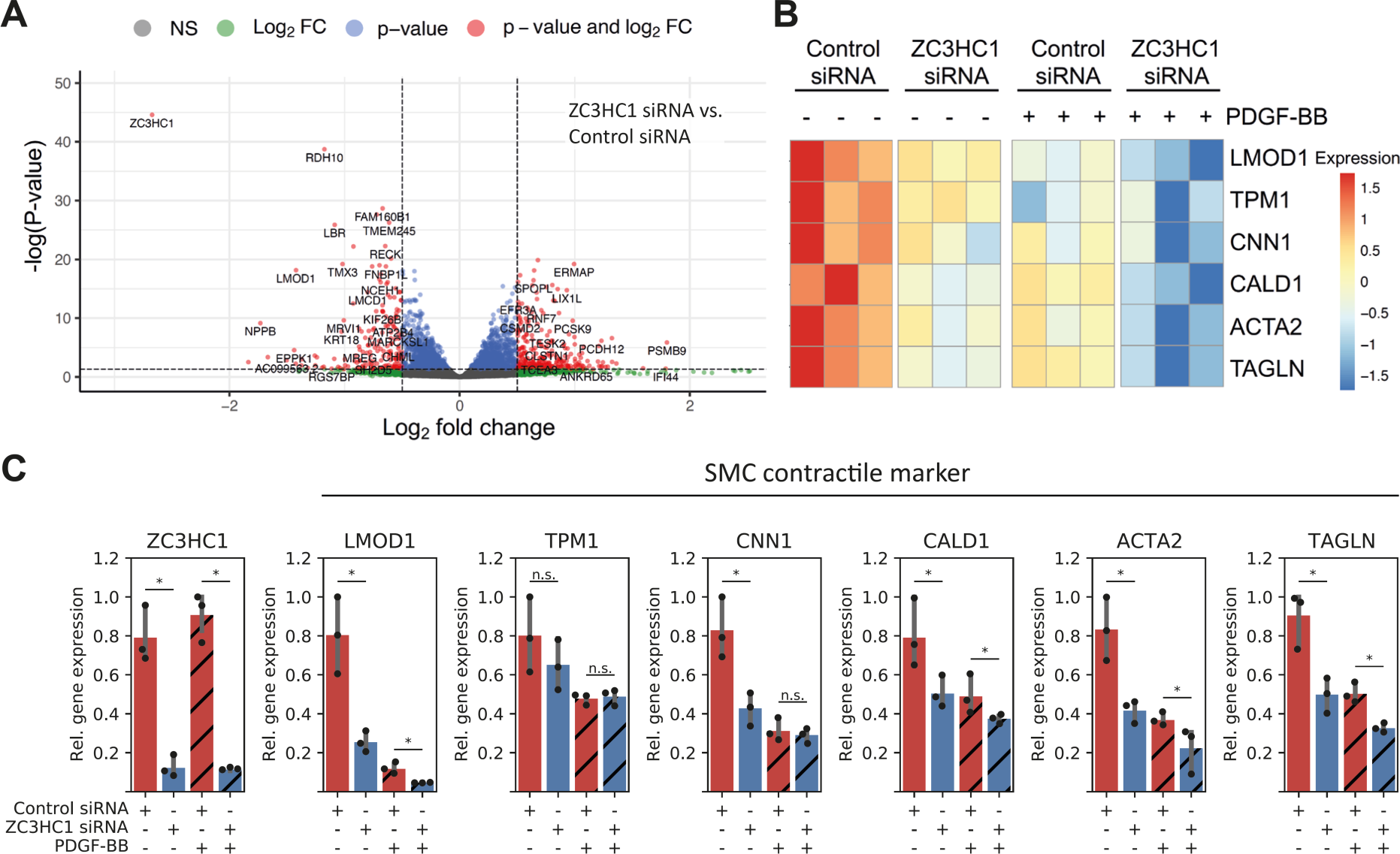
Transcription profiling of human SMCs after *ZC3HC1* downregulation. (**A**) Volcano plot of the expression profiles of differentially expressed genes in SMCs transfected with siRNA against *ZC3HC1* (*ZC3HC1* siRNA) (n=4) and control siRNA (n=6) using PDGF-BB treatment as co-variate. The red data points represent the differentially expressed genes with statistical significance, whereas the gray data points indicate genes without disturbed gene expression. The vertical dashed lines correspond to a 0.5-fold-change in gene expression (up or down), and the horizontal dashed line represents the adjusted p-value for each gene. (**B**) Heat map showing contractile SMC marker genes that were differentially expressed upon knockdown of *ZC3HC1* (from one experimental batch; i.e. 3 replicates per condition) in the presence or absence of PDGF-BB. (**C**) qPCR validation of the expression of the contractile SMC marker genes. Values are shown as mean ± s.d.; * p<0.05; n.s., not significant.

## Network analysis of differentially expressed genes

Subsequent network analysis of differentially expressed genes (p_adj_<0.05 and abs(log_2_(FC))>0.4) using the STRING database^27^ revealed that the *CCNB1* gene encoding cyclin B1 is a central hub for a variety of gene clusters (**Figure 3A-B, Supplementary Figure S2 and Supplementary Table 6-8**), whereas *ZC3HC1* interacts only with *CCNB1* (STRING score=0.607). Therefore, network analysis indicated that modulation of *ZC3HC1*/NIPA predominantly induces transcriptional changes through cyclin B1. Additional enrichment analysis of gene clusters highlighted several biological processes that were apparently modulated by the knockdown of *ZC3HC1,* including protein ubiquitination, muscle contraction/ cytoskeleton organization, and cell division (**Figure 3B**). To test whether *CCNB1* is modulated by lower levels of *ZC3HC1* in SMCs, we performed co-expression and Western blot analyses. Analysis of gene expression in aortic SMCs of 151 individuals showed a positive correlation (R=0.59; p=1.4 × 10^-15^) between *ZC3HC1* and *CCNB1* at the RNA level (**Figure 3C**, top panel). Similarly, qPCR showed that siRNA mediated *ZC3HC1* downregulation reduced *CCNB1* expression in SMCs (**Figure 3C**, top right). At the protein level, however, a lack of NIPA resulted in the intracellular accumulation of cyclin B1 (**Figure 3C** bottom panel), as previously described ^13^. To explain changes in the expression of SMC contractile marker genes (**Figure 2C** and **Figure 3A/3B**), we analyzed protein levels of the transcription factor serum response factor (SRF) which is critical for maintaining the contractile phenotype of SMCs ^37^. Indeed, the level of native SRF is significantly impaired in the presence of PDGF-BB in *ZC3HC1* deficient SMCs (**Figure 3D**), indicating a *ZC3HC1-CCNB1-SRF* axis regulating the expression of SMC contractile marker genes (e.g., *CNN1*). On the protein level, however, no significant changes could be observed for *CNN1* or *LMOD1* (**Figure 3E**), demonstrating the minor effects of short term *ZC3HC1* silencing in SMCs.

**Figure 3.**
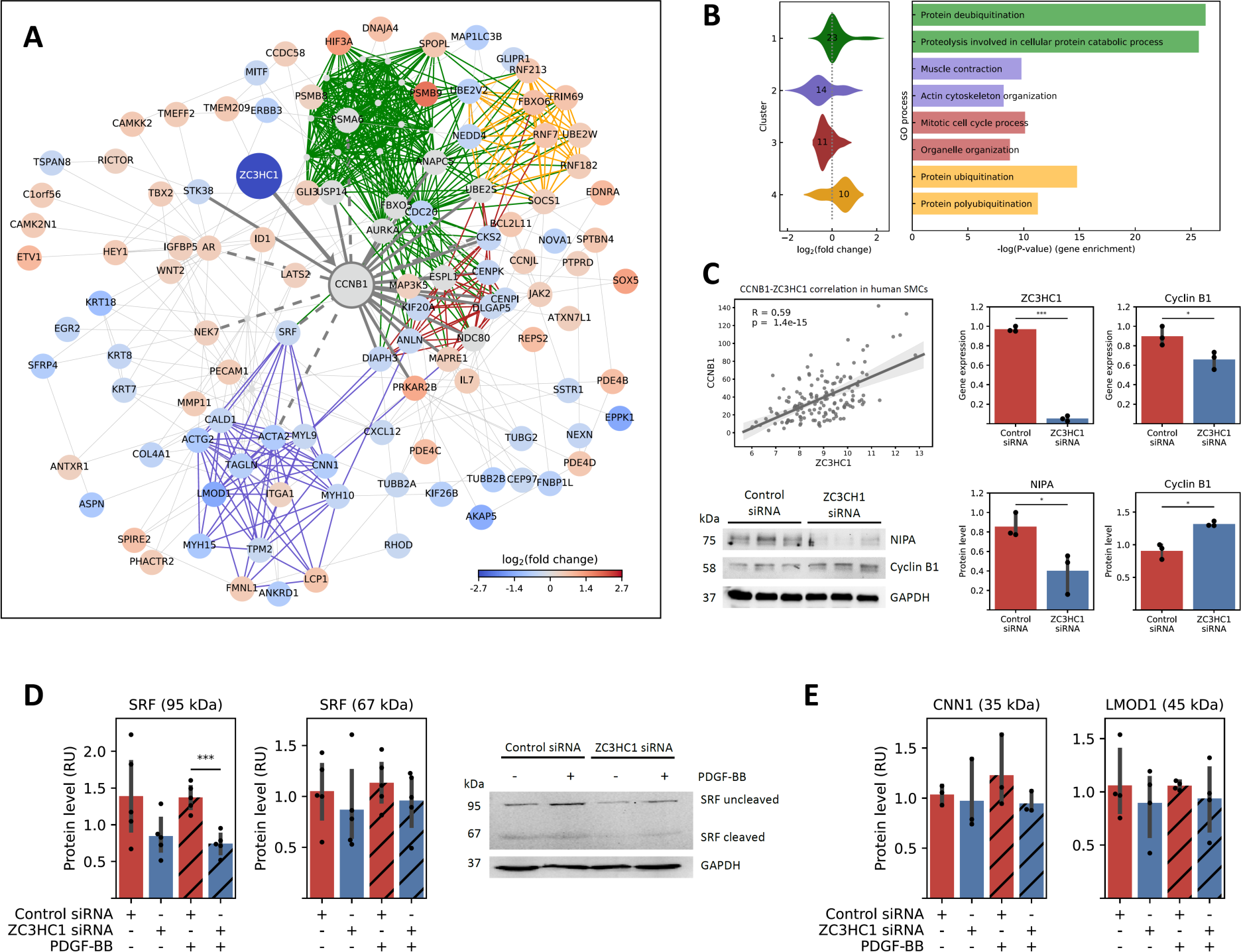
Gene interaction network of *ZC3HC1*/CCNB1 in SMCs and accumulation of cyclin B1 by *ZC3HC1* knockdown. (**A**) Effect of *ZC3HC1* knockdown on the cyclin B1 (CCNB1) subnetwork of differentially expressed genes (adjusted p-value <0.05 and abs(log_2_(fold-change))>0.4) derived from the STRING database. The edges represent the combined STRING scores (>0.4, median=0.9) derived from co-expression, experimental, database, and text mining scores. The complete gene interaction network is shown in **Supplementary Figure S2**. The fold changes in gene expression on the log_2_-scale are depicted by red and blue spheres. For example, SMC contractile marker genes (see purple cluster 2 in A and B) show lower gene expression (see also Figure 2B and 3B). Gray spheres indicate genes not differing in expression but interacting with CCNB1 or neighboring genes according to the STRING database. Gray bold lines indicate direct interactions with CCNB1 with medium to high confidence (scores >0.5), and dashed lines indicate direct interactions with lower confidence (score=0.4–0.5). Colored lines represent gene clusters. (**B**) Distribution of log_2_(fold changes) in gene expression for each cluster and enriched biological processes derived from Gene Ontology (GO). Numbers in the violin plot indicate the number of genes in the cluster. (**C**) Co-expression analysis in 151 human aortic SMCs preparations showed a positive correlation between *ZC3HC1* and *CCNB1* gene expression, which was confirmed by transient knockdown of *ZC3HC1* and qPCR (right top corner). By contrast, Western blot showing that *ZC3HC1* knockdown results in the accumulation of cyclin B1 protein in SMCs. (**D**) Western blot analysis of SRF (serum response factor) confirmed a significant reduction of SRF protein level in ZC3HC1-deficient SMCs in the presence of PDGF-BB. (**E**) For CNN1 and LMOD1, only a small trend towards lower protein levels has been observed in ZC3HC1 knockout SMCs. Values of bar plots are shown as mean ± s.d.; * p<0.05; *** p<0.001.

## Proteomic analysis of SMC proteins

In addition to RNAseq, we subjected cell lysates from SMCs carrying the T allele (n=7) or C allele (n=6) or from *ZC3HC1* knockdown SMCs versus control (n=4 per condition) to proteomic analyses. As expected for SMCs (four replicates) transiently treated with *ZC3HC1* specific siRNA compared to control (scrambled siRNA), changes at protein level were only subtle and not significant after adjusting for multiple testing (**Supplementary Figure S3** and **S4**). However, proteomic analysis of SMCs carrying the rs11556924-T allele (ZC3-T) versus SMCs carrying the rs11556924-T allele (ZC3-C) revealed that only twelve proteins show significantly lower protein levels in ZC3-T SMCs (**Supplementary Figure S5** and **Supplementary Table 9**). These proteins (e.g., platelet-derived growth factor receptor beta) are seemingly involved in PDGF signaling (Enrichr^38^: BioPlanet 2019; p-value=0.003; q-value=0.08; odds ratio=363.3) and cell chemotaxis (Enrichr: Gene ontology biological process; p-value=0.0008; q-value=0.06; odds ratio=59.5).

### Phenotyping of *Zc3hc1*^-/-^ mice

The murine homolog of the human *ZC3HC1* gene is ubiquitously expressed in various tissues (**Supplementary Figure S6A-B**). X-Gal staining confirmed that transgenic mice harbored the *LacZ* gene (**Supplementary Figure S6C**) with similar results observed at the DNA level using agarose gel electrophorese for *LoxP* sites (**Figure 4A**). Phenotypically, the birth rate was lower for *Zc3hc1***^-/-^**than for WT and heterozygous animals. In general, Zc3hc1^-/-^ mice display reduced birth rate (males [8%]; females [11.5%]) and are infertile. In addition, knockout of *Zc3hc1* had a significant impact on body weight (**Figure 4B**, left), with lifetime body weight being significantly lower for *Zc3hc1***^-/-^** than for WT (23±2g vs. 27±5 g; p<0.0001 at 44 weeks) (**Supplementary Figure S6D**). In addition, a few *Zc3hc1***^-/-^** mice exhibited shorter snouts than their WT littermates (**Figure 4 B**, right). Despite the effect of *Zc3hc1* knockout on body weight, life span was similar in *Zc3hc1***^-/-^**and WT mice (**Figure 4B**, bottom).

**Figure 4.**
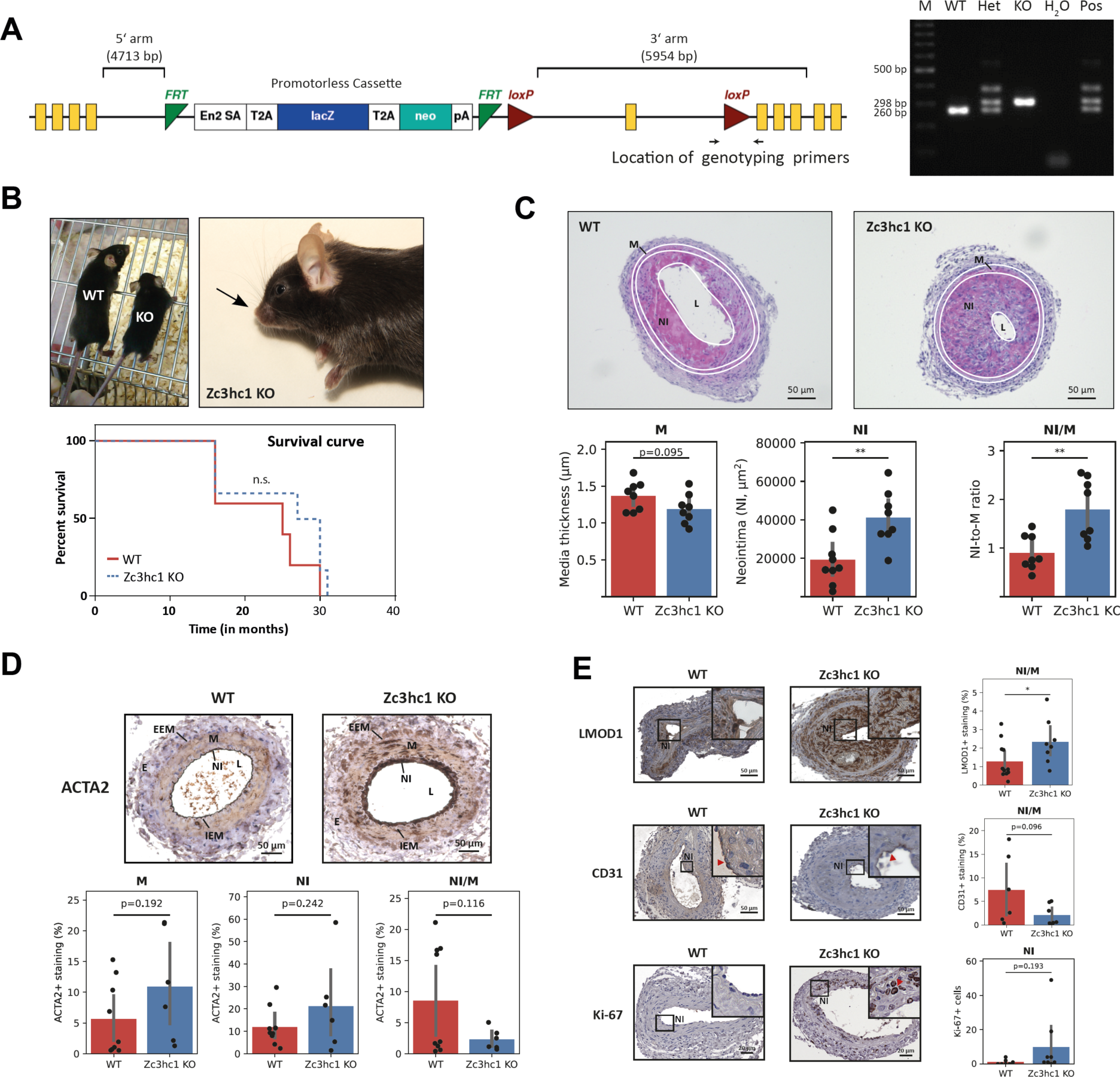
Generation and characteristics of *Zc3hc1^-/-^* (KO) mice. (**A**) Targeting vector used to generate *Zc3hc1^-/-^* mice (left) and genotyping of mice by PCR (right) (M: 100 bp marker, wild-type mice (WT or *Zc3hc1*^+/+^); Het: Heterozygous mice (*Zc3hc1*^+/-^); *Zc3hc1* knockout mice (KO or *Zc3hc1*^-/-^); H_2_O; Pos: Positive control represented by Het mice (*Zc3hc1*^+/-^). Location of genotyping primers. Primer pair targeting the intron between exons 5 and 6 of *Zc3hc1*, which contains a *loxP* site (left panel) in the targeted allele. The sizes of the PCR products were 298 bp in KO (*Zc3hc1*^-/-^) and 260 bp in WT (*Zc3hc1*^+/+^) mice, with Het mice (*Zc3hc1*^+/-^) possessing both alleles. (**B**) Decreased body weight (left) and abnormally short snout (indicated by arrow) (right) in KO mice. The survival curve showing no statistically significant difference between KO (n=6) and WT (n=5) mice. **C)** Representative hematoxylin & eosin-stained femoral artery sections (top), and quantification of media thickness (M), area of neointima formation (NI) and neointima-to-media ratios (NI/M) (bottom) in female WT (n=8) and female KO (n=8) mice. Scale bars, 50 µm. (**D**) ACTA2-positive cells predominantly reside in the media (M) and in the neointima (NI) in both WT and Zc3hc1 KO mice (E, Externa; EEM, external elastic membrane; IEM, internal elastic membrane). (**E**) Tissue slides were stained with anti-LMOD1 (SMC marker), anti-CD31 (Endothelial cell marker), or anti-Ki-67 (proliferation maker) antibody. Primary antibodies were visualized using a horseradish peroxidase system and DAB staining solution. A detailed description can be found in the main text. Values of the bar plots are shown as mean ± s.d.; * p<0.05; ** p<0.01; n.s., not significant.

### NIPA deficiency leads to increase neointima formation as a response to injury in mice

Because of the effects of *ZC3HC1* on the *in vitro* migration and proliferation of human SMCs, the role of *Zc3hc1* on neointima formation was assessed *in vivo* using vascular injury model. Neointima formation was induced in *Zc3hc1^-/-^*and WT mice by wire injury of the femoral artery, and the extent of neointima formation was assessed after 14 days. Neointima formation was approximately 2-fold greater in KO than in WT mice (p=0.005), accompanied by an increase in intima-to-media ratio (p=0.003) (**Figure 4C**). No major significant differences were observed in the vessel circumference between Zc3hc1*^-/-^* and littermate controls after injury (73.9 ± 12.2 µm^2^ vs. 61.5 ± 20.6 µm^2^; p=0.155). Although ACTA2 gene expression was significantly decreased in human *ZC3HC1* knockdown SMC, histochemical analysis of femoral arteries sections suggests no difference in the abundance of ACTA2-positive cells; neither in the media nor neointima between KO and WT mice (**Figure 4D)**. The SMC specific marker LMOD1 points towards a significant higher SMC content in the neointima (NI) in *Zc3hc1* KO mice versus WT mice (**Figure 4E**). In line, we found less CD31-positive cells (endothelial cells) in the NI in KO than in WT mice and slightly more Ki-67 positive cells being in the G2 and M phase (**Figure 4E**), although the results were not statistically significant. These results indicate that vascular remodeling after injury is predominantly driven by the migration of medial SMCs.

### Murine *Zc3hc1* knockout SMCs are contractile, less proliferative, and more migratory

Based on the findings from human SMCs, we isolated primary aortic SMCs from mice (female and male) and tested whether NIPA deficiency in murine primary aortic SMCs modifies migration and proliferation. Compared with WT SMCs, the migration of *Zc3hc1***^-/-^**SMCs was significantly enhanced at 12 h (p=0.04), but not at later time points (**Figure 5A**). Furthermore, the proliferation of *Zc3hc1***^-/-^** SMCs was significantly diminished than that of WT SMCs (**Figure 5B**). This effect is enhanced in PDGF-BB-induced proliferation. Because NIPA has been shown to interact with cyclin B1^13^, we compared the expression of cyclin B1 protein in NIPA-deficient and WT murine SMCs. In line with decreased proliferation, *Zc3hc1***^-/-^** SMCs show significantly reduced cyclin B1 levels as compared to WT SMCs (p=0.043) (**Figure 5C**). Further, *Zc3hc1*^-/-^ SMCs exhibit a contractile signature reflected by enhanced gene expression of almost all SMC contractile marker genes (**Figure 5D**) and higher protein levels of *Lmod1* and *Cnn1* (**Figure 5E**). This contractile SMC phenotype was observed in both female (**Figure 5**) and male SMCs lacking *Zc3hc1* (**Supplementary Figure S8**).

**Figure 5.**
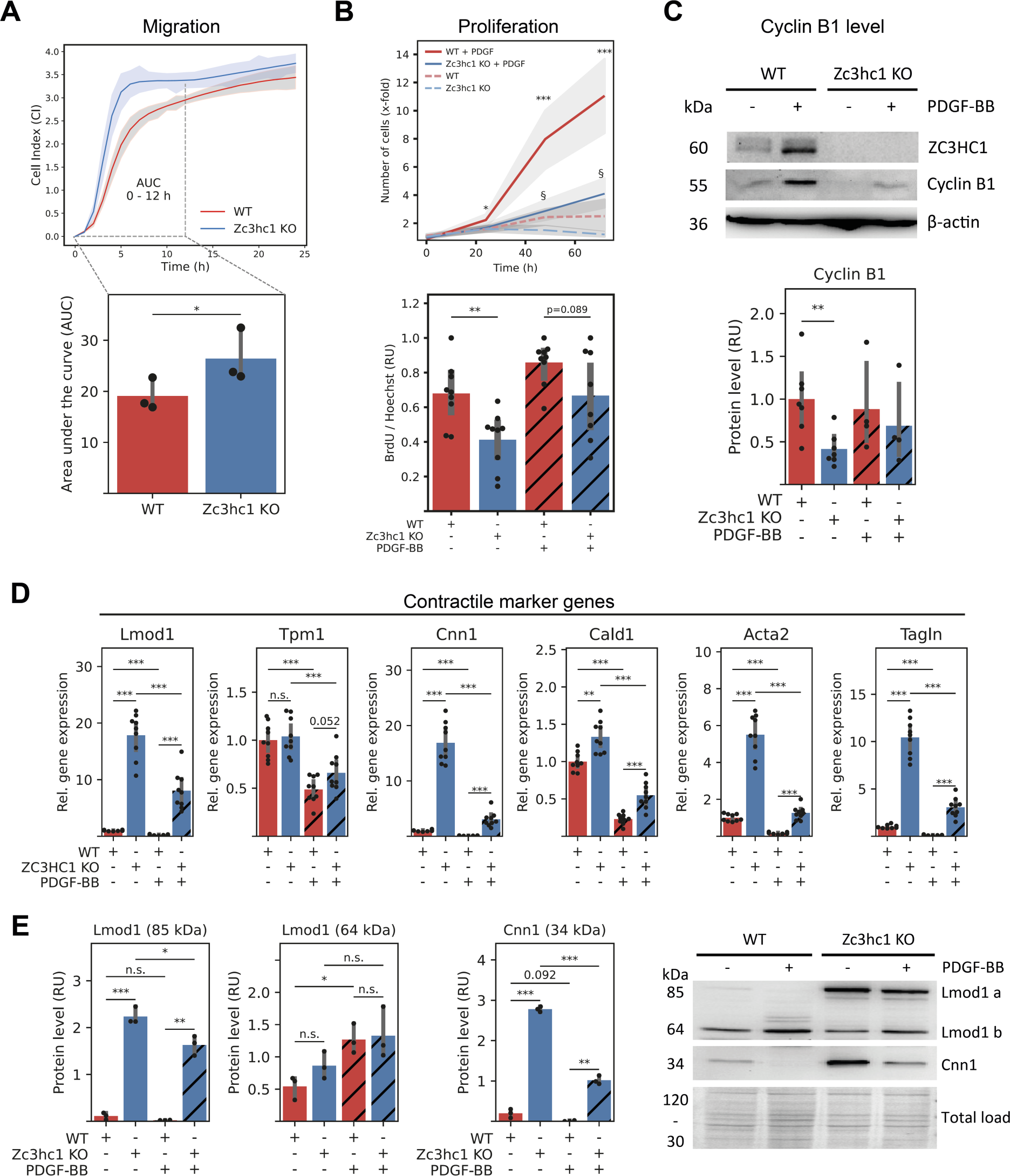
Knockout (KO) of *Zc3hc1* in murine aortic SMCs. SMCs isolated from female *Zc3hc1* knockout (KO) mice show elevated migration in the first 12 h (**A**) but less proliferation (**B**) compared with SMCs isolated from wild-type (WT) mice. (**C**) Western blotting of NIPA/ZC3HC1 protein confirming the knockout of *Zc3hc1* in SMCs, resulting in reduction of cyclin B1 protein. (**D**) Relative mRNA levels are shown for contractile SMC marker genes. Values were normalized to WT without PDGF-BB (mean = 1). (**E)** Western blot analyses of Lmod1 and Cnn1. A representative Western blot is shown right (Total load = amido black staining). The translucent error bands (A-B) represent one standard deviation. Values of bar plots are shown as mean ± s.d.; * p<0.05; ** p<0.01; *** p<0.001, p-values below 0.1 are shown as numbers else not significant (n.s.)

### NIPA and cyclin B1 are located at similar sites during mitosis

To further understand the mechanistic link between *ZC3HC1*/NIPA, cyclin B1 and cell migration/proliferation we captured the location of both proteins during the cell cycle in murine SMCs using confocal microscopy. During interphase, NIPA is predominantly located in the nucleus (NC), but also appears to be present in the cytoplasm (CP) where it colocalizes to some degree with microtubules **(Figure 6**). During mitosis, NIPA seems to be located either in fragmented nuclear membranes^39,40^ in the prophase or at the cleavage furrow during anaphase and telophase. On the contrary, cyclin B1 is mainly present outside the NC in small amounts (**Figure 7**). The level of cyclin B1 starts to rise during G2 phase and cyclin B1 accumulates in the NC during the G2 to M phase transition^41^. In the prometaphase and metaphase, cyclin B1 appears to overlap with components of the spindle apparatus (e.g. centrosomes^42^ and microtubules^43^). Similar to NIPA, cyclin B1 appears to be located directly at the contractile actomyosin ring^44^ during telophase and cytokinesis. Of note, a similar colocalization of NIPA and cyclin B1 during late mitosis and cytokinesis was also observed in primary human SMCs (**Supplementary Figure S9**). No obvious structural differences were observed between mitotic WT SMCs and *Zc3hc1* KO SMCs for cyclin B1. However, there is a tendency towards more plasma membrane blebbing in *Zc3hc1* KO SMCs during (pro)metaphase (see **Figure 7B**). To estimate the proportion of SMCs in different cell cycle states, we counted SMCs in G2 phase, G2-M transition, and M phase (**Figure 7C**). A total of 144 images were analyzed using an in-house Python script that uses staining intensity and location (NC vs. CP) of cyclin B1 to assign the different phases. In accordance with lower proliferation rates of murine *Zc3hc1^-/-^* SMCs, the number of nuclei is lower in *Zc3hc1^-/-^*SMCs than in WT SMC (673 ± 131 vs. 1143 ± 131), as is the percentage of SMCs undergoing G2 (4.3 ± 1.2% vs. 18.4 ± 3.8%), G2-M transition (0.8 ± 0.4% vs. 3.3 ± 1.0%), or mitosis (0.6 ± 0.4% vs. 1.9 ± 0.6%). Interestingly, the proportion of SMCs undergoing mitosis versus SMCs in G2/G2-M transition phase is higher in *Zc3hc1^-/-^* SMCs than in WT SMCs, providing weak evidence that the absence of *Zc3hc1*/NIPA results in a longer mitosis phase.

**Figure 6.**
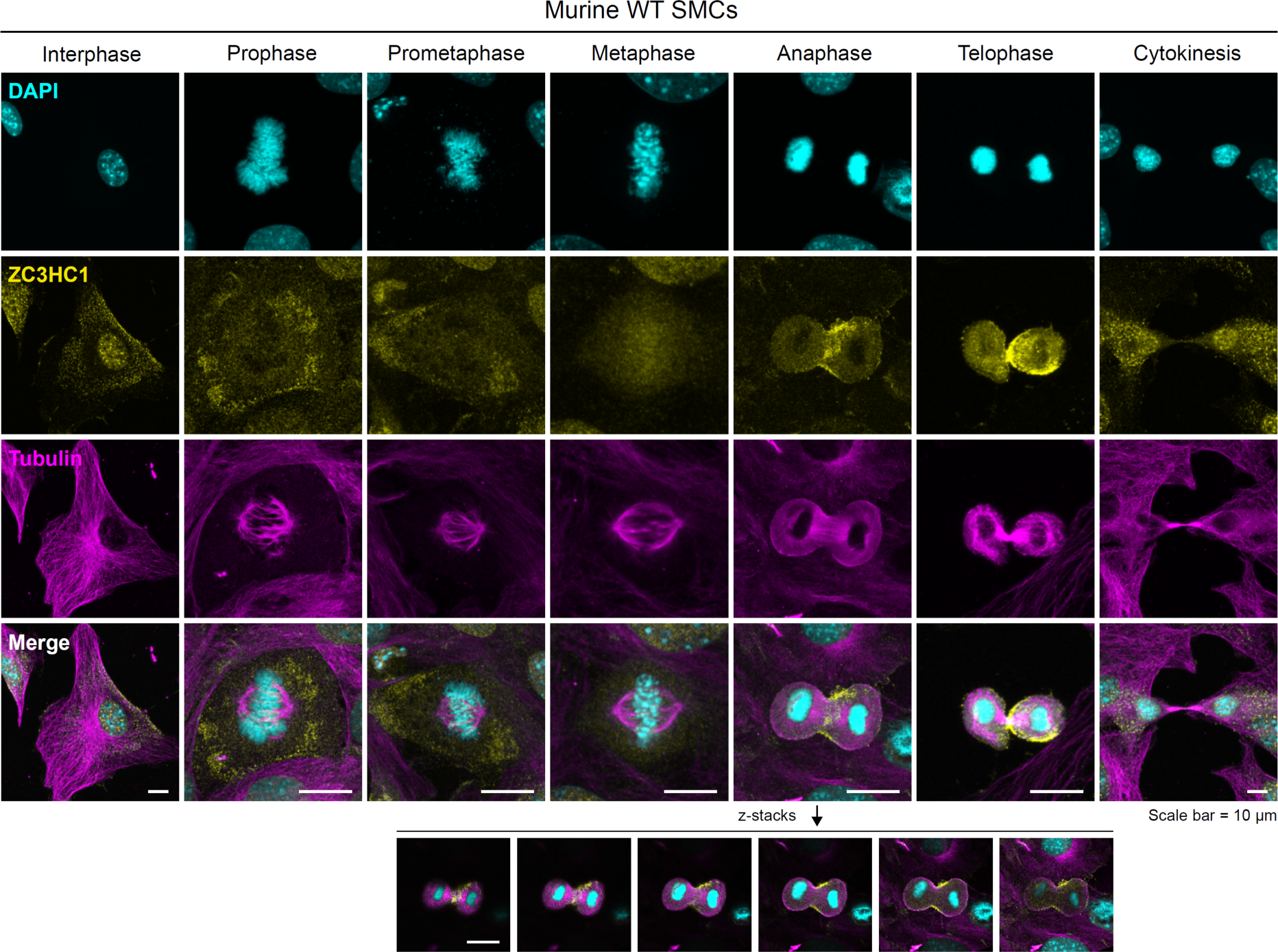
Localization of Zc3hc1 protein in murine aortic SMCs during the cell cycle. Pseudo-colored confocal immunofluorescence images are maximum intensity projections of z-stacks (∼1 µm distance). PDGF-BB-treated murine wild-type (WT) SMCs were stained with DAPI (cyan), anti-ZC3HC1 (yellow), anti-tubulin (magenta). During interphase, the *Zc3hc1* protein *(Zc3)* is predominantly located in the nucleus but is also colocalized to some extent with microtubules. During the initial stage of mitosis (prophase/prometaphase), Zc3 appears to be located at fragmented nuclear membranes^39^. Later during the metaphase, anaphase, and early telophase, Zc3 is predominantly situated in the cleavage furrow and contractile ring. The location of Zc3 during the anaphase is highlighted in the lower panel.

**Figure 7.**
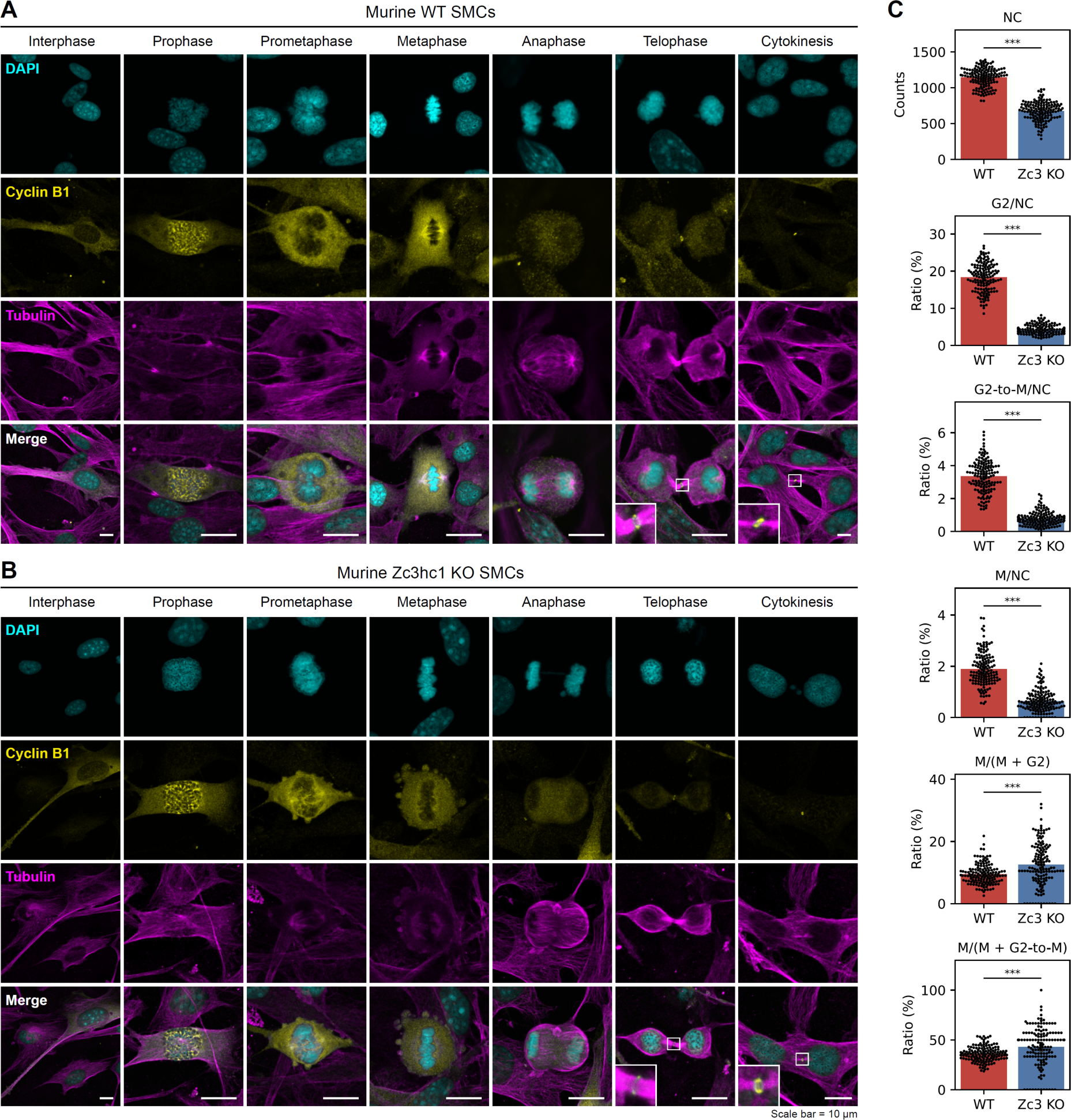
Localization of cyclin B1 in murine aortic SMCs during the cell cycle. Pseudo-colored maximum intensity projections of confocal immunofluorescence z-stacks (∼1 µm distance) show PDGF-BB treated (**A**) wild-type (WT) and (**B**) *Zc3hc1* (Zc3) knockout (KO) murine SMCs stained with DAPI (cyan), anti-cyclin B1 (yellow), anti-tubulin (magenta), and respective secondary antibodies. During interphase, cyclin B1 resides in the cytoplasm at low levels. At early prophase, cyclin B1 is accumulated in the nucleus where it is involved in the disassembly of nuclear pore complexes^45^ and depolymerization of nuclear lamin filaments through phosphorylation^46^. During prometaphase and metaphase, cyclin B1 overlaps partly with components of the spindle apparatus (centrosomes^42^ or spindle assembly checkpoints^43^). In the late stage of mitosis (telophase) and during cytokinesis, cyclin B1 appears to be directly located at the contractile actomyosin ring^44^ (enlarged sections). (**C**) shows the number of nuclei (NC) of murine SMCs (WT vs. Zc3 KO) as well as the number of murine SMCs in G2 phase, in the G2 to M phase transition (G2-to-M), and SMCs in mitosis (M). A total of 144 images per genotype from 3 replicates taken with a 10x objective were analyzed semi-automatically using an in-house Python script (available upon request). Values of bar plots are shown as mean ± s.d.; one dot represents one image *** p<0.001.

## Discussion

Restenosis is one of the main clinical complications in patients who undergo coronary artery revascularization ^47,48^. SMCs regulate the arterial contractile tonus and blood pressure and constitute the key cell type during atherosclerotic plaque formation and in response to revascularization procedures ^49–51^. SMCs migrate from the media into the intima of the vessels, followed by alterations in their phenotype in response to their new microenvironment ^49–51^. This study assessed the role of *ZC3HC1* on the migration, proliferation and neointima formation of SMCs. The ubiquitously expressed *ZC3HC1* gene encodes the cell cycle protein NIPA, which has been found first to be associated with CAD in several independent GWASs ^2–8^. The CAD-associated rs11556924-C/T SNP in *ZC3HC1* is functional, leading to an amino acid change from arginine to histidine (p.Arg363His) ^4,7,9^. Analysis of the publicly available GTEx dataset eQTL (V8) showed that the genetic variant rs11556924-T results in reduced *ZC3HC1* gene expression in the heart (atrial appendage and left ventricle) and skin samples. Because eQTL effects are often cell type-specific, we tested whether an eQTL was present in aortic SMCs from 151 heart transplant donors^19^, finding that SMCs from donors carrying the rs11556924-T allele have lower *ZC3HC1* expression and migrate faster than SMCs from donors carrying the rs11556924-C allele (**Figure 1B – C**). Of note, the migration phenotype observed here might not be linked to the slight decrease of *ZC3HC1* but to the presence of the protein variant. We previously reported^52^ the absence of a significant association between *ZC3HC1* expression and rs11556924 in blood samples^9^, monocytes/macrophages^35^ and aortic endothelial cells^36^, suggesting that the observed regulatory impact of the variant at the *ZC3HC1* locus is specific to SMCs.

To further evaluate and find more evidence supporting the role of *ZC3HC1* in SMC biology and its putative contribution to vascular remodeling, we used a commercially available human SMC line and siRNA knocked down the expression of ZC3HC1 to mimic the lower expression observed in rs11556924-T allele carriers. Similarly, siRNA-mediated KD of *ZC3HC1* in human SMCs resulted in increased cell migration during the first 12 h compared with controls. Consistent with previous results ^9^, PDGF-BB-induced proliferation of *ZC3HC1* KD SMCs was greater than control SMCs at 72 h, but not at earlier time points (e.g., 24 h) (see **Figure 1D – F**). This may explain the lack of a significant correlation between the rs11556924-T allele and cell proliferation in our study at 24 h. Further, downregulation of *ZC3HC1* or its protein NIPA in primary human SMCs led to increased cyclin B1 protein levels, in agreement with previous findings ^7,13^ (**Figure 3C**).

To assess the molecular mechanisms involved in *ZC3HC1* modulation, we performed transcriptome analysis, followed by analyses of gene/protein interaction networks and pathway enrichment *in vitro*. The fact that a reduction of *ZC3HC1* mRNA results in a transition of SMCs from a contractile/quiescent phenotype to a proliferative, synthetic and migratory phenotype^49–51,53^ was reflected in reduced gene expression of canonical SMC contractile marker genes such as *alpha-smooth muscle actin* (*ACTA2*), *calponin* (*CNN1*), *transgelin* (*TAGLN*), and *leiomodin 1* (*LMOD1*) ^49–51,53–55^. In addition to the reduction in gene expression of SMC contractile marker genes, our pathway analysis revealed that an impaired level of *ZC3HC1* seems to promote a more active, non-quiescent state of SMCs as genes highly expressed in *ZC3HC1* KD SMC are enriched for extracellular matrix (ECM) organization, cytokine-mediated signaling, and G protein-coupled receptor signaling (**Supplementary Figure S2**). However, the direct molecular consequences of modulating ZC3HC1 are still vague. Besides the involvement of cyclin B1, a central regulatory node in our gene interaction network, in regulating cell proliferation (discussed later), protein ubiquitination may be an important factor. Many genes involved in protein ubiquitination (e.g., *FBX06*, *RNF7*, *RNF213*, and *RNF182*) and proteasome complex formation (e.g., *PSMB8* and *PSMB9*) are upregulated in *ZC3HC1* KD SMCs, probably to compensate for the deficiency of NIPA as part of the SCF complex. For example, *FBX06* (see **Figure 3A**, yellow cluster) is a component of the SCF-type E3 ubiquitin ligase complex^56^. SCF complexes, in general, have been found to control cell proliferation through ubiquitin-mediated degradation of critical regulators, including cell cycle proteins (e.g., cyclins) or transcription factors (e.g., β-catenin) ^57^. Therefore, NIPA deficiency leading to an increased expression of genes involved in protein ubiquitination (see **Figure 3A**, yellow cluster) may trigger an orchestrated gene regulation accompanied by ubiquitin-mediated degradation of specific transcription factors such as serum response factor (SRF). Indeed, we observed lower levels of SRF (**Figure 3A** and **Figure 3D**) due to *ZC3HC1* knockdown. The transcription factor SRF targets about one-quarter of differentially expressed genes, including *ZC3HC1* itself and *CCNB1* explaining their positive correlation at mRNA levels (**Figure 3C**); moreover SRF targets all canonical SMC contractile marker genes analyzed ^58,59^.

As *ZC3HC1* KD induced transcriptional changes are rather small, with only 60 genes being up- or downregulated ≥2-fold, many of its effects may occur at the protein level, e.g., cyclin B1 accumulation (as shown) and/ or posttranslational protein modifications. For example, the Cdk1-cyclin B1 complex phosphorylates the protein Mcl-1, a regulator of apoptosis, thereby inducing its degradation ^60^. We performed mass spectrometry analysis to evaluate the global impact of ZC3HC1 KD at the protein level. However, we only detected significant differences in SMCs from 151 heart transplant donors with different rs11556924 genotypes (T vs. C), but not in ZC3HC1 KD SMCs (**Supplementary Figures S3 – S5**); suggesting that global changes in protein levels may only occur over time (e.g., several weeks) following modulation of ZC3HC1/NIPA.

Besides its known function as component of the SCF-type E3 ubiquitin ligase complex, and direct Cyclin B1 interactor, it has been recently postulated that the zinc finger protein ZC3HC1/NIPA resides in nuclear baskets of vertebrate nuclear envelopes ^15^. As nuclear basket protein, ZC3HC1/NIPA presumably recruits TRP encoding coiled-coil proteins^15^ which are components of the nuclear pore complex and required for the export of mRNA (and proteins) ^15,16^. Moreover, these ZC3HC1-dependent TPR polypeptides^15^ are also involved in chromatin compartmentalization in the interphase and mitotic spindle checkpoint signaling during mitosis ^16^. Therefore, *ZC3HC1* KD might also affect nuclear pore function, thereby modulating SMC states and cell cycle processes. These findings indicate that *ZC3HC* modulation initiates various steps involving protein ubiquitination and degradation of specific factors that normally maintain the contractile phenotype in human SMCs.

To determine whether these *in vitro* findings could be extended to an *in vivo* model, we investigated the effects of complete NIPA KO on vascular remodeling in a mouse model. Like human SMCs, SMCs from these mice migrated faster compared with SMCs derived from their WT littermates. The increased migratory activity of murine SMCs may explain the greater degree of injury-induced neointima formation in *Zc3hc1***^-/-^**compared to WT mice (**Figure 4C**), since the amount of LMOD1 protein (a typical SMC marker) is significantly higher in *Zc3hc1* KO mice (**Figure 4E**, upper panel).

In contrast to human *ZC3HC1* KD SMCs, the complete absence of NIPA in murine SMCs resulted in less cell divisions (**Figure 5B**), decreased cyclin B1 levels and increased expression of SMC contractile marker genes (**Figure 5D-E**), which might be explained by impaired nuclear pore function as well as co-expression of *ZC3HC1* and *CCNB1* (**Figure 3C**), regulated in part by SRF. The lower proliferation rate may also explain why Zc3hc1^-/-^ mice are significantly smaller compared to WT mice (**Supplementary Figure S6**). To understand why the lack of *ZC3HC1* results in lower cell proliferation, we examined the location and putative role of NIPA during interphase and mitosis (M) using confocal immunofluorescence microscopy in murine (**Figure 6 and 7**) and human SMCs (**Supplementary Figure S9**). Based on these findings, we found that NIPA and cyclin B1 are located at similar sites during mitosis, especially in the nucleus during G2 to M transition and at the cleavage furrow/contractile ring during late phases of mitosis. Therefore, we postulate that NIPA is involved in regulating cyclin B1 during several steps of cell division. Cyclin B1 forms with Cdk the M phase promoting factor (MPF) which is a key component in the control of cell cycle progression^13^ and phosphorylates several proteins involved in mitosis, e.g., microtubules^61^ and even NIPA at Ser-395^7,62^. Its level starts to rise during G2 phase and cyclin B1 accumulates in the nucleus (NC) during the G2 to M transition^41^ due to inactivation of NIPA by phosphorylation of Ser-354 and Ser-359 ^63^. Inside the NC, the MPF (cyclin B1 *+* Cdk) starts mitotic onset by phosphorylating nuclear membrane components such as nuclear pore complexes^45^ and lamin filaments^46^ leading to disassembly of the nuclear envelope. As part of the nuclear core complex^15^, NIPA could tightly regulate this process. Similarly, NIPA could regulate processes at the cleavage furrow, as cyclin B1-Cdk1 is required for anillin phosphorylation^44^ and anillin is important for cleavage furrow organisaton^64^, cytokinesis, and bleb assembly during mitosis ^65,66^. The latter would explain why we observe more membrane blebbing in *Zc3hc1^-/-^* SMCs during mitosis^67,68^. Notably, gene expression of *ANLN* (encoding anillin protein) is also regulated by SRF and correlates with *CCNB1* expression (STRING score=0.891; see **Figure 3A**, red cluster), indicating co-regulatory mechanisms.

The interaction between NIPA and cyclin B1 was also demonstrated using a genome-editing approach in the pseudo-diploid colon carcinoma cell line DLD-1 ^7^. The change in amino acid (p.Arg363His; rs11556924-T) was found to alter cyclin B1 dynamics, presumably resulting from enhanced NIPA phosphorylation at Ser-395. Phosphorylated NIPA cannot form the CSF^NIPA^ complex which normally ubiquitinates cyclin B1 ^13^. The lack of cyclin B1 ubiquitination accelerates its nuclear accumulation, reducing the time required to complete mitosis ^7^. On contrary, a complete knockout of NIPA in murine SMCs appears to slow down mitosis, as the percentage of SMCs in mitosis versus SMCs in G2/G2-M transition was higher in *Zc3hc1^-/-^*SMCs than for WT SMCs (**Figure 7C**). Moreover, cyclin B1-Cdk1 promotes mitotic progression indirectly through upregulation of the anaphase-promoting complex/cyclosome (APC/C)^69^, and a lower basal level of cyclin B1 could impede this process. This may also explain the trend toward more Ki-67 staining, a proliferation marker, in the neointima of femoral artery isolated from *Zc3hc1^-/-^* mice (**Figure 4E**), because Ki-67 is directly involved in multiple steps of mitosis ^70^. However, further experiments are needed to confirm this assumption. Because fewer Zc3hc1^-/-^ SMCs were in G2 phase compared to WT SMCs, it is plausible that Zc3hc1^-/-^ SMCs have higher migratory ability, since cells in late S or G2 phase do not migrate ^71,72^. Overall, it appears that the effect of modulation of ZC3HC1/NIPA is biphasic and there might be a tipping point. While lower levels of NIPA promote SMC proliferation presumably due to cyclin B1 accumulation, a complete loss hampers cell division probably because of lower cyclin B1 levels, less nuclear pore integrity, and impaired mitosis (anaphase/ telophase) or cytokinesis.

Taken together, our *Zc3hc1* KO mouse model suggests that the complete lack of NIPA results in an exacerbated injury-induced neointima formation most likely through SMC migration from the media towards neointima. Cell cycle or mitosis arrested SMCs in the neointima might recruit more medial SMCs to migrate owing to insufficient healing of the injured vessel. Given the association of SCF complexes with genome stability and cancer pathogenesis^69^, it is also conceivable that loss of *Zc3hc1* in mice results in a few SMCs with cancerous signatures that drive the neointima formation. However, we were unable to demonstrate this in cell culture. In patients with *ZC3HC1* alterations (lower level and/or missense variants), the increased neointima formation could be due to increased SMC migration and enhanced cell proliferation within the neointima. Since a higher level of cyclin B1 is associated with more proliferation and enhanced neointima formation^73^, and inhibition of this increase protects against neointima formation ^74,75^. Additional studies are required to determine the exact molecular mechanisms at which level *ZC3HC1* affects cellular proliferation, migration and neointima formation, for example, using SMC/EC-lineage tracing mouse models and/or pharmacological approaches. For this, using a humanized knock-in model is required.

Finally, it remains elusive why the T allele in *ZC3HC1* lowers the risk of CAD ^4,6,9,10^, but increases the risk of carotid IMT as demonstrated recently in 502 patients with rheumatoid arthritis ^11^. One reason could be the bivalent role of vascular SMCs as their biological function in atherosclerosis have opposite effects depending on the stage of the lesions (early vs. advanced stage). At early stages of the formation of atherosclerotic lesions, migratory and proliferative SMCs are known to play a detrimental role in atherosclerosis ^49,76–78^. Therefore, anti-proliferative therapies were previously proposed for atherosclerosis ^79,80^.

On contrary, the role of SMCs in advanced lesions is thought to be beneficial by stabilizing the plaque, which may result in an asymptomatic progression of atherosclerosis. Interestingly, two working groups observed that SMCs derived from advanced plaques are less proliferative. Consequently, they assumed that enhancement and not inhibition of SMC proliferation might be beneficial for plaque stability in advanced lesions ^78,80^, thereby reducing the risk of major adverse cardiovascular event outcomes such as MI. Moreover, the T allele also lowers the risk for blood pressure (**Figure 1A**), a major risk factor in CAD ^81,82^, probably due to its impact on NIPA gene expression/phosphorylation and SMC contractile genes.

Taken together, our integrative analyses highlighted the functional role of *ZC3HC1* as a neointima formation-associated gene. This might offer clues into potentially targetable SMC-mediated disease mechanisms. However, the role of NIPA on cell cycle regulation and SMC biology seems to be more complex as previously anticipated (**Figure 8**), considering the location of NIPA during cell cycle (SCF complex, nuclear pore complex, and cleavage furrow), different phosphorylation sites, the tight regulation between NIPA and cyclin B1 (NIPA phosphorylation by cyclin B1-Cdk1 vs. cyclin B1 ubiquitination by NIPA), and the co-expression of *ZC3HC1* and *CCNB1*.

**Figure 8:**
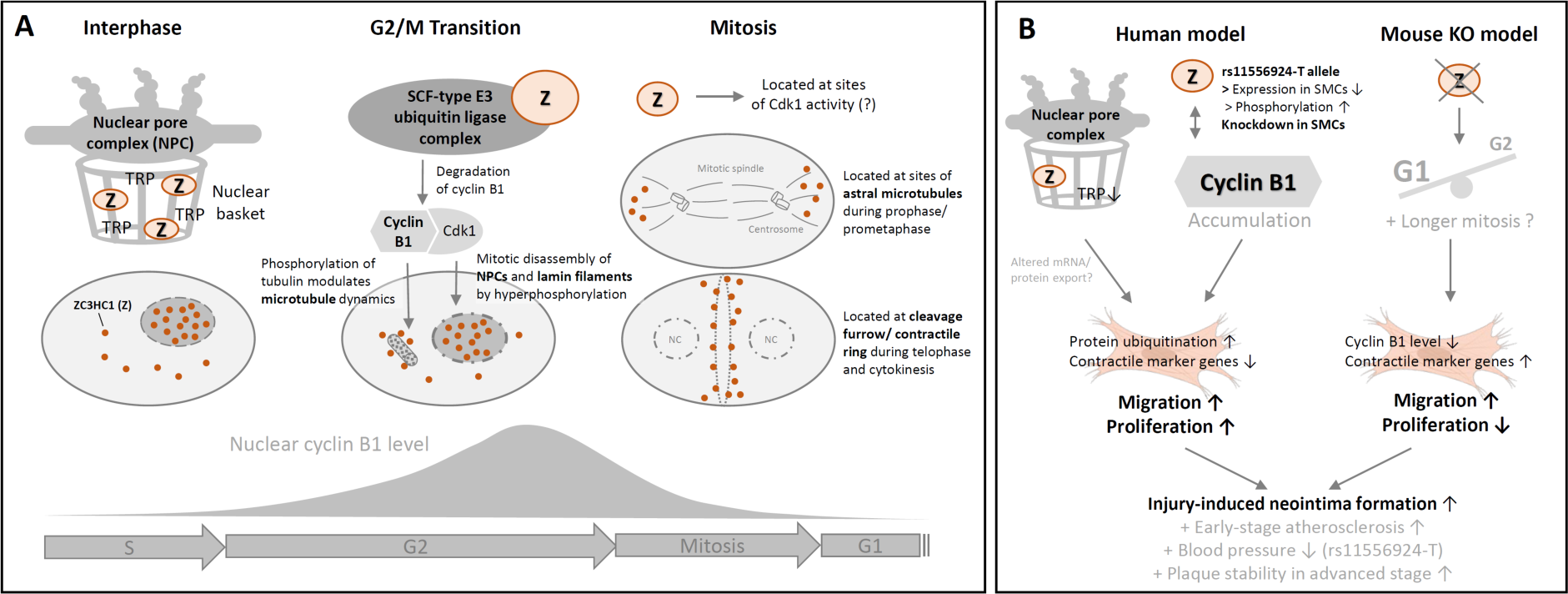
Diagram illustrating the proposed mechanisms of ZC3HC1/NIPA modulation based on the main findings of this study and literature review. (A) NIPA is part of the nuclear pore complex^15^ and the SCF-type E3 ubiquitin ligase complex^7,13^. During mitosis, NIPA appears to be located at sites of cyclin B1/ cyclin-dependent kinase 1 (Cdk1) activity^41^, including nuclear pore complexes^45^, astral microtubules^61,83^, and cleavage furrow^44,64^. **(B)** Under normal (healthy) conditions NIPA activates the degradation of cyclin B1 during G1, S, and early G2 phase. However, *ZC3HC1* modulation by reducing the amount of NIPA or increasing NIPA phosphorylation^7,9^, is accompanied by cyclin B1 accumulation and increased SMC migration and proliferation, which might also be mediated by dysfunctional nucleocytoplasmic transport of mRNAs and proteins through less recruitment of TRP coiled-coil polypeptides^15^. This in turn leads to injury-induced neointima formation and progression of early-stage atherosclerosis ^49,76–78^. The rs11556924-T is associated with lower blood pressure (see **Figure 1A**) and increased SMC proliferation in the late stage of atherosclerosis may contribute to plaque stability ^78,80^, resulting in an asymptomatic progression of this disease. However, the complete lack of Zc3hc1 in murine SMCs results in reduced levels of cyclin B1 and cell cycle arrest, presumably in the M/G1 phase. Therefore, neointima formation in Zc3hc1 knockout (KO) mice is most likely a consequence of SMC migration rather than cell proliferation. Illustrations of cells are from BioRender.com.

Our findings provide strong evidence that lower *ZC3HC1* gene expression in the presence of the rs11556924-T allele or through genetic manipulation in SMCs increased cell migration. Consequently, we postulate that more migration/proliferation of SMCs increases the risk of early lesion progression and intima-media thickening or neointima formation in humans but may have an advantage in advanced lesions by stabilizing the plaque.

## Limitations

The number of studies showing an association between *ZC3HC1* and intima-media thickening in inflammatory conditions such as rheumatoid arthritis is limited. Also the number of patients (n= 500) included is very low. Therefore, more human cohorts with more cases of IMT are desirable. Furthermore, we cannot rule out any sex bias for the development of IMT, because only female mice were included in our study. However, we did not see any sex-stratified effects in our human dataset or datasets from the UK Biobank (**Supplementary Table 2**). Considering the proposed multiple functions of *ZC3HC1*/NIPA, a complete knockout model should be extended towards an SMC/EC-lineage tracing mouse model in which Zc3hc1 is only transiently or moderately modulated in a cell-specific manner. Moreover, knockin mouse will be the ideal model to study the exact role of the rs11556924-T/C (p.363Arg/His).

## Author contributions

Redouane Aherrahrou, Tobias Reinberger, Heribert Schunkert, Thorsten Kessler, Jeanette Erdmann and Zouhair Aherrahrou designed the project. Redouane Aherrahrou, Tobias Reinberger, Jeanette Erdmann and Zouhair Aherrahrou contributed to the text of the main manuscript. Redouane Aherrahrou, Tobias Reinberger, Jaafar Al-Hasani, Julia Werner, Miriam Otto, Sandra Wrobel, Maren Behrensen, and Zouhair Aherrahrou performed the characterization experiments and generated data for most of the figures and tables. Maria Loreto Munoz-Venegas, Mete Civelek and Thorsten Kessler participated in data analysis and lookup. All authors contributed to the final article.

## Sources of Funding

This work was supported by the German Federal Ministry of Education and Research (BMBF) in the context of the German Centre for Cardiovascular Research (FKZ 81Z0700108, FKZ81X2700133); by Fondation Leducq (18CVD02, PlaqOmics) and its Junior Investigator Award (to R.A., and T.R); and by the Deutsche Forschungsgemeinschaft (DFG, German Research Foundation) under Germany’s Excellence Strategy – EXC 22167-390884018 to HB and as part of the collaborative research centers SFB 1123 (B02, to T.K. and H.S.), TRR 267 (B06, to H.S.). This study was also supported by an American Heart Association Postdoctoral Fellowship (18POST33990046, to RA), National Institutes of Health grant (R21 HL135230, to MC), the University of Eastern Finland (FI22857339, to RA) and by the German Federal Ministry of Education and Research (BMBF) in the context of the DZHK-Säule-B projects (FKZ 81X2700121). In addition, this work was supported by the Corona Foundation as part of the Junior Research Group Translational Cardiovascular Genomics (S199/10070/2017).

## Acknowledgments

The authors thank Maren Behrensen, Sandra Wrobel, and Lisa Paurat for their technical support. We thank Jan Wenzel (Institute of Pharmacology and Toxicology, University of Lübeck) for technical support with confocal microscopy. We also thank the members of the Erdmann, Schunkert, and Civelek laboratories for their feedback and discussions.

## Conflict of interest

The authors declare no conflict of interest.

## Notes

### Competing Interest Statement

The authors have declared no competing interest.

### Summary of Updates

Here we updated the in vivo findings using our knockout (Zc3hc1-/-) mice by showing enhanced neointima formation in response to arterial injury and faster SMCs migration ability. However, complete loss of Zc3hc1 led to a significant reduction in SMC proliferation and lower cyclin B1 protein level. Moreover, immunostaining and confocal microscopy demonstrated, for the first time, that ZC3HC1 and Cyclin B1 were located at the cleavage furrow during mitotic progression of SMCs.

