## Supplementary material for "Deficiency of *ZC3HC1* modulates vascular smooth muscle cell phenotype and increases neointima formation": Suppl_Figures_S-S9

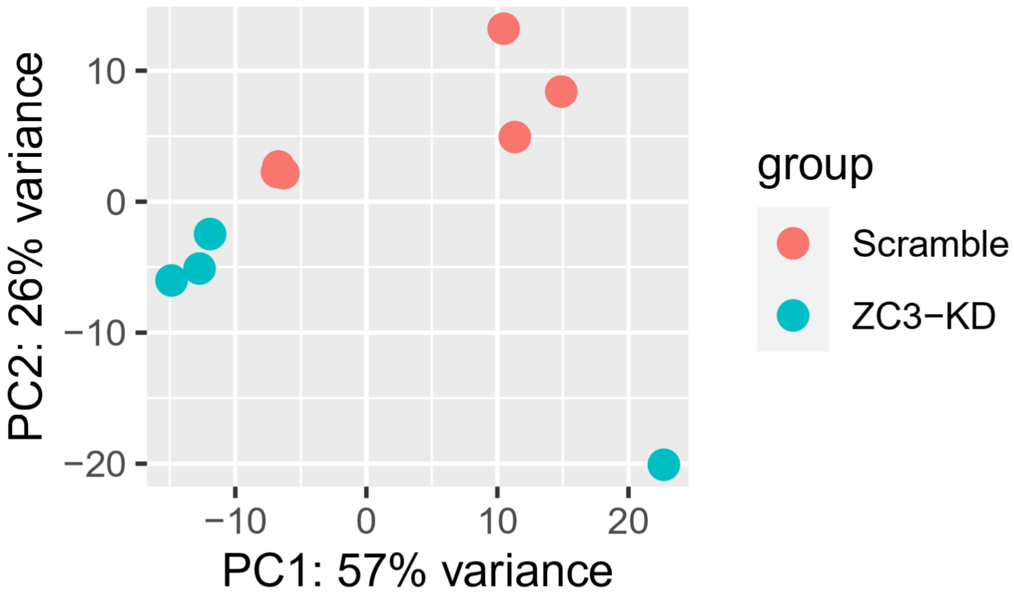


**Figure S1:** Principal component analysis (PCA) of gene expression in SMC transfected with siRNA against ZC3HC1 (ZC3-KD; n=4) and control siRNA transfected cells (Scrambled; n=6).

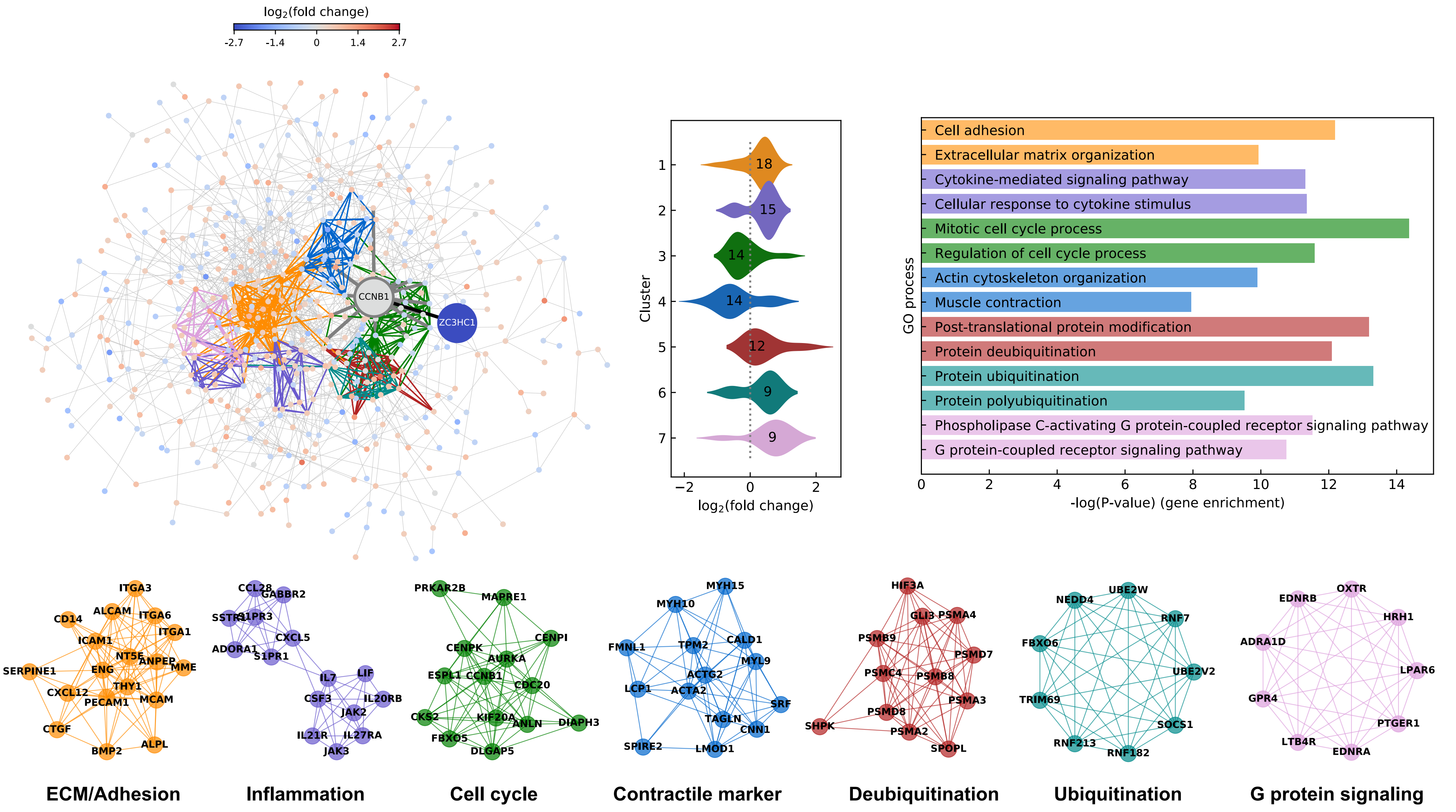


**Figure S2**: Complete STRING interaction network of ZC3HC1 KD induced differentially expressed genes (padj<0.05) with abs(log2(fold change))>0.4. ZC3HC1 downregulation led to CCNB1 encoded cyclin B1, a central hub gene in the network. STRING scores (>0.4, median=0.9) were derived from co-expression score, experimental score, database score, and text mining score. The fold change in gene expression on the log2-scale is depicted by red and blue spheres, respectively. Gray spheres show no difference in gene expression but interactions with CCNB1 or neighboring genes according to the STRING database. Colored lines represent gene clusters. The right panel shows the distribution of log2(fold change) in gene expression for each cluster and enriched biological processes derived from Gene Ontology (GO). Numbers in the violin plot display the number of genes in this cluster.

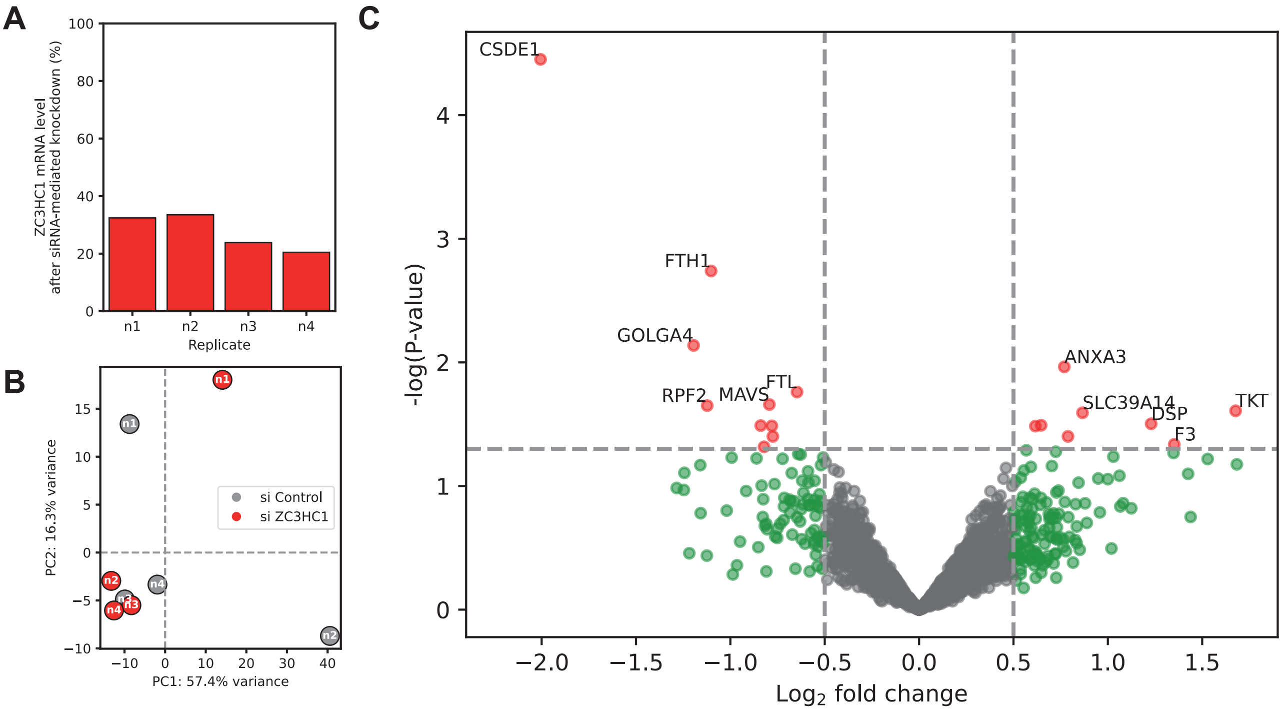


**Figure S3: (A)** Residual ZC3HC1 mRNA levels for four independent technical replicates compared to respective negative controls (scrambled siRNA) following siRNA-mediated knockdown in human aorta smooth muscle cells treated with 5 ng/ml PDGF-BB for 24 h. **(B)** Principal component anaylsis (PCA) for eight samples. **(C)** Volcano plot with non-adjusted p-values. Only for CSDE1 (Cold shock domain-containing protein E1), the adjusted p-values (p_adj = 0.063) is close to statistical significance after adjusting for multiple testing using the Benjamini-Hochburg method as implemented in the R LIMMA package.

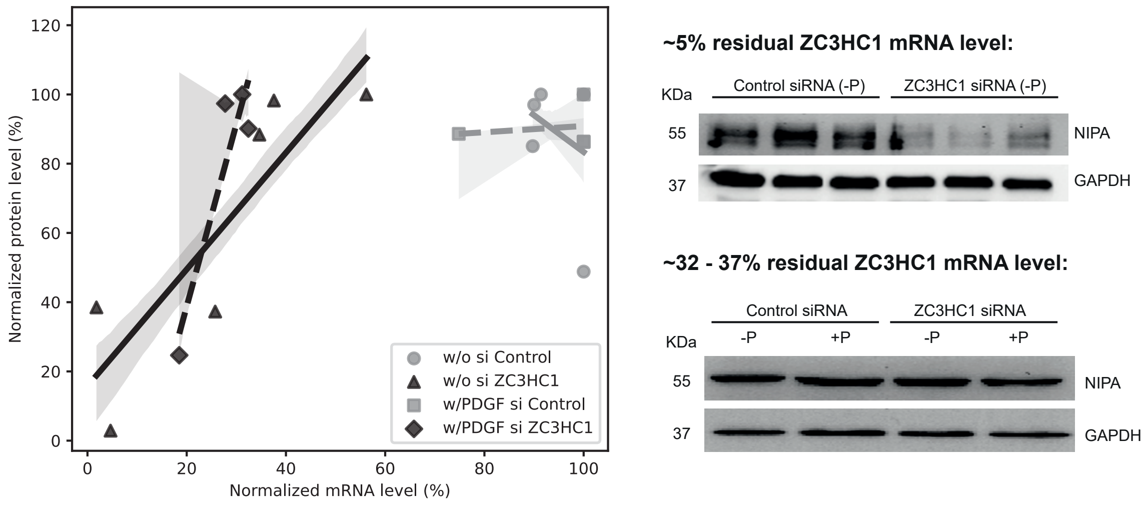


**Figure S4:** Relation between ZC3HC1 mRNA levels and ZC3HC1/NIPA protein levels. Less than ~25% residual ZC3HC1 mRNA levels are required after siRNA-mediated knockdown to achieve notable changes at protein level. Human aorta smooth muscle cells were treated with 5 ng/ml PDGF-BB (w/ or +P) for 24 h or without PDGF-BB (w/o or -P). See Material and Method section in the main text for more details.

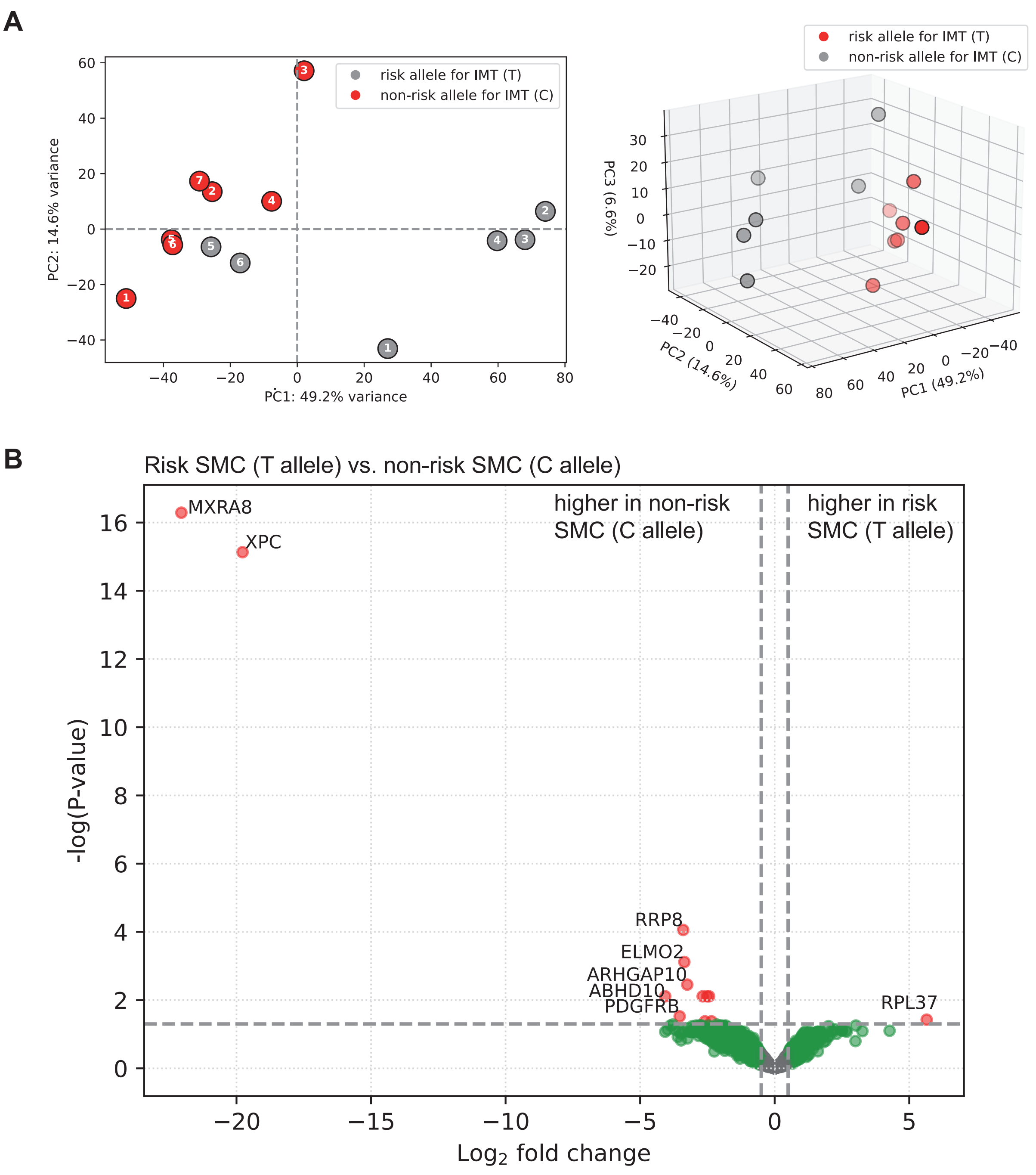


**Figure S5: Proteomic analysis of human vascular smooth muscle cells carrying the T allele or C allele. (A)** Principal component analysis (PCA) for six homozygous SMC carrying the risk allele T (SMC-ZC3-T) and seven SMC carrying the non-risk allele C (SMC-ZC3-C). **(B)** Volcano plot of the differential protein level analysis using the R LIMMA package comparing risk SMC-ZC3-T and non-risk SMC-ZC3-C. P-values represent adjusted p-values after multiple test correction using the Benjamini-Hochburg method as implemented in the R LIMMA package. The proteins MXRA8 (Matrix Remodeling Associated 8) and XPC (XPC Complex Subunit, DNA Damage Recognition and Repair Factor) were undetectable in samples derived from risk SMC-ZC3-T. Gene annotations can be found in supplementary table ST9.

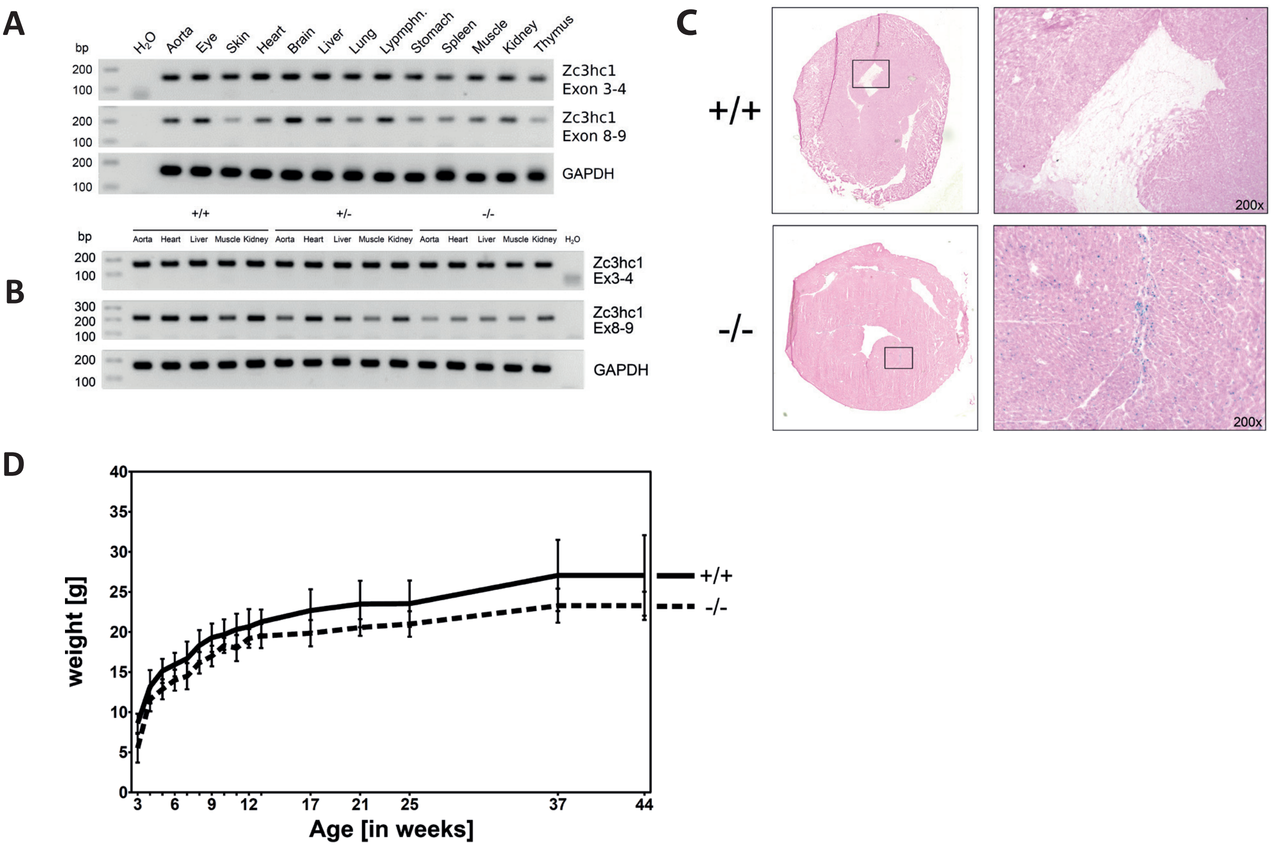


**Figure S6:** Establishing and phenotyping the *Zc3hc1*-knockout mouse model. **A)** To obtain an expression profile for Zc3hc1, mRNA from various organs of a wildtype mouse was isolated, transcribed to cDNA and used for a PCR assay for *Zc3hc1*. Both primer pairs used, one between exon 3 and 4, the other between exon 8 and 9 show bands in all organs, at the respective expected product size for each pair, indicating a ubiquitous expression of *Zc3hc1*-mRNA. GAPDH was use das a loading control. **B)** Full-length mRNA is expressed also in transgene animals. cDNA from different organs was tested using primers targeting a sequence before (exon 3 and 4) and after (exon 8 & 9) the inserted cassette, respectively; both products are seen in all organs from all three genotypes. GAPDH was used as loading control. $H_2O$: water control; bp: base pairs. **C)** Functional expression of the inserted lacZ-Gene. X-Gal staining shows expression of beta-Galactosidase, the enzyme encoded by the gene lacZ, in HET and KO but not in WT animals. Heart cryosections from animals of all three genotypes were stained with X-Gal, which releases a blue chromogen if metabolized by beta-Galactosidase. Left: overview of the heart section. Right: detailed view of the rectangular areas marked in the heart sections. D) *Zc3hc1*-KO mice (n=7) have a significantly ($p < 0.0001$, paired t-test) lower body weight compared to WT mice (n=16). Initially, mice were weighed weekly (age 3 to 13 weeks), followed by weight controls at the depicted time points.

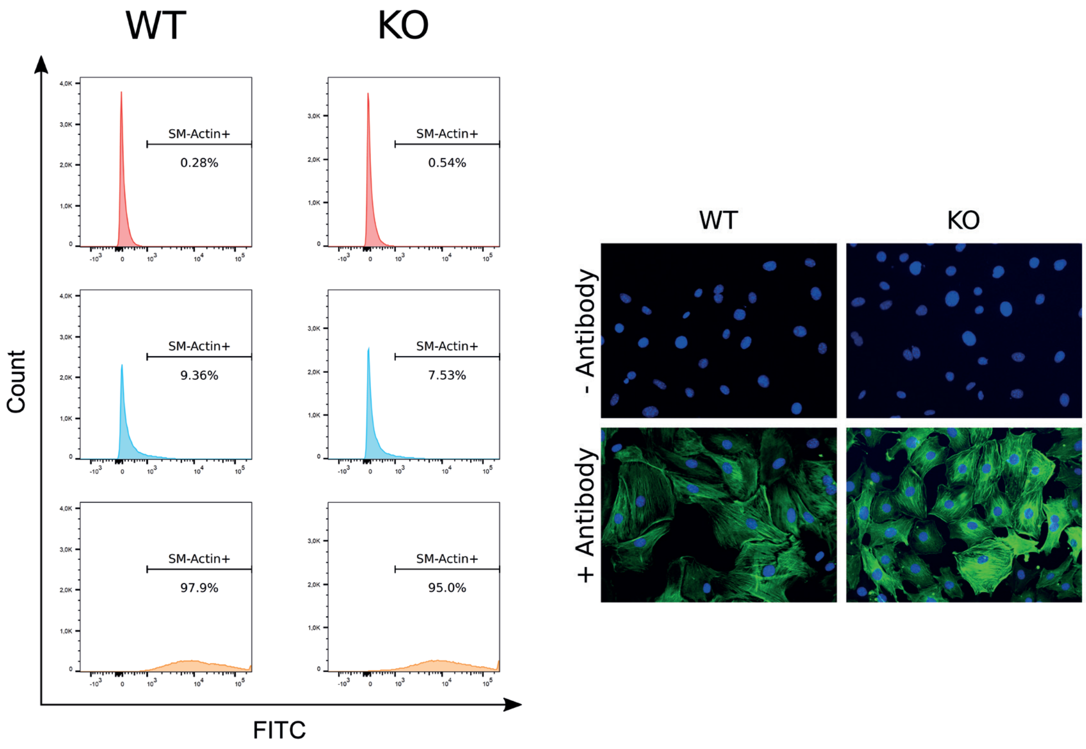


**Figure S7:** Quantification of SMC marker alpha-SMA. Cells were analysed by Flow Cytometry. Top row: unstained; Middle row: Isotype Control; Bottom row: anti-alpha-SMA antibody. Characterization of SMCs using the specific marker alpha-SMA. Shown are the overlay images of cells stained with the FITC-labeled anti-alpha-SMA antibody (green) and DAPI to visualize nuclei (blue). Top row: controls (without primary antibody). Bottom row: the typical actin pattern is visible in both WT and KO SMCs.

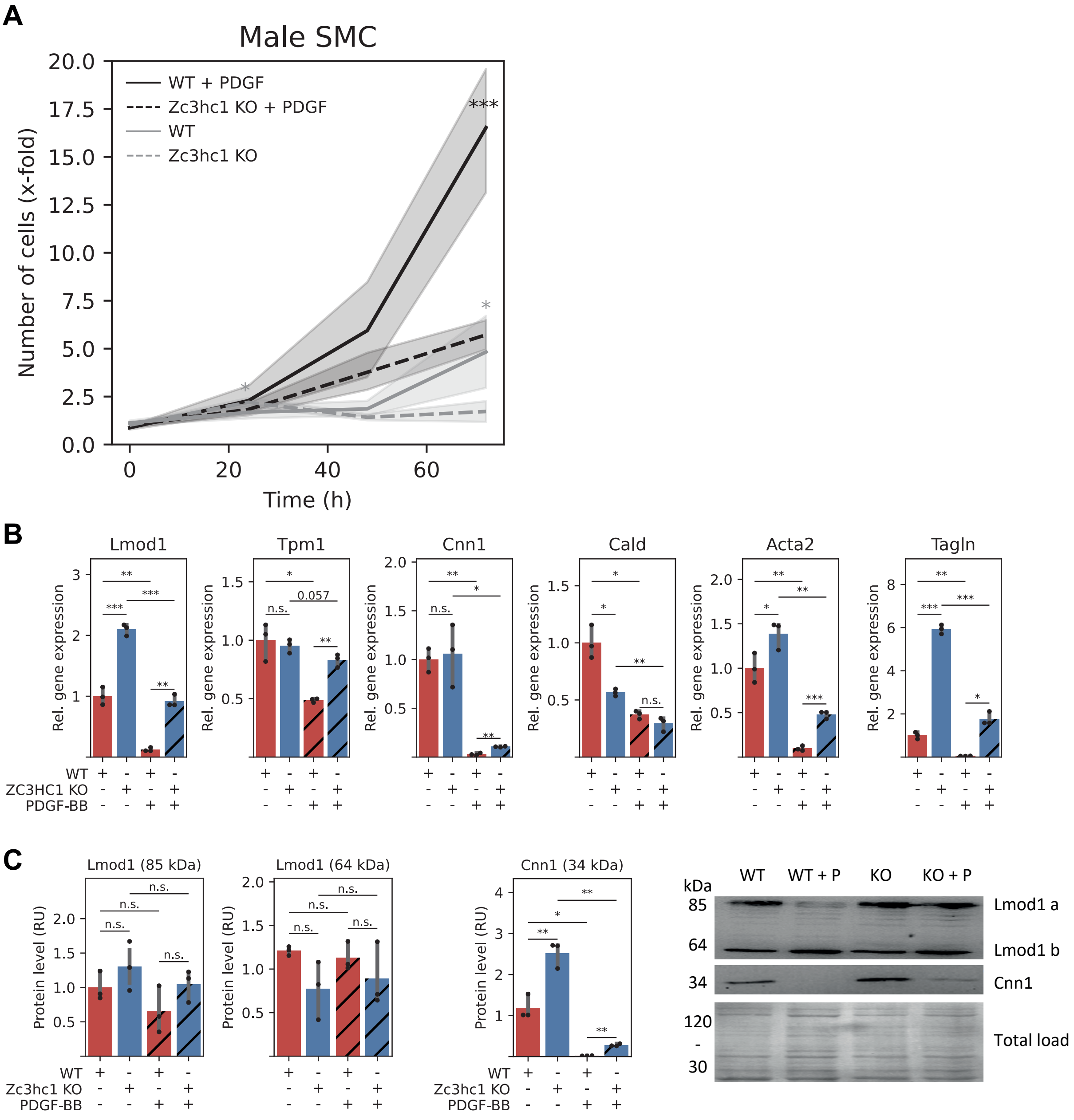


**Figure S8:** Phenotypic analysis of aortic smooth muscle cells isolated from male *Zc3hc1*-knockout (KO) mice and wildtype (WT) mice. Like female *Zc3hc1*-KO SMCs (see Fig. 5), *Zc3hc1* deficiency results in lower proliferation **(A)** and significant higher level of Lmod1, Tpm1, CNN1, Acta2, and Tagln in presence of PDGF-BB as demonstrated by quantitative PCR **(B)** and in part by Western blot **(C)**. Values of bar plots are shown as mean ± s.d.; * p<0.05; ** p<0.01; *** p<0.001. Data were analyzed using unpaired Student's t-test.

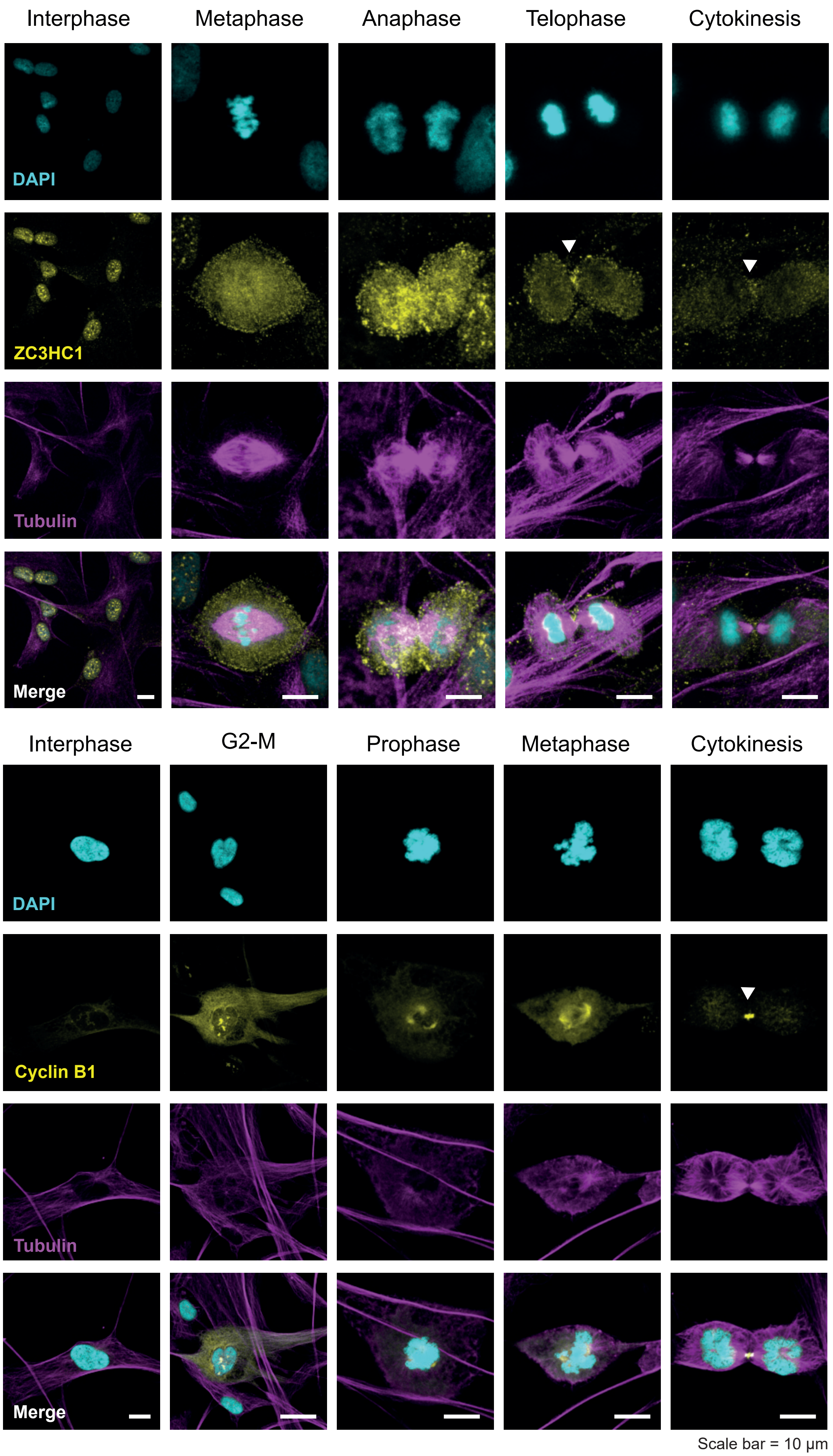


**Figure S9. Localization of Zc3hc1 protein and cyclin B1 in human aortic SMCs during the cell cycle.** Pseudo-colored confocal immunofluorescence images are maximum intensity projections of z-stacks (~1 µm distance). Human SMCs were stained with DAPI (cyan), anti-ZC3HC1 or anti-cyclin B1 (yellow), and anti-Tubulin (magenta). During telophase and cytokinesis, ZC3HC1 and cyclin B1 appear to be located in proximity to the contractile ring (white arrows).
